# Polygenicity and epistasis underlie fitness-proximal traits in the *Caenorhabditis elegans* multiparental experimental evolution (CeMEE) panel

**DOI:** 10.1101/120865

**Authors:** Luke M. Noble, Ivo Chelo, Thiago Guzella, Bruno Afonso, David D. Riccardi, Patrick Ammerman, Adel Dayarian, Sara Carvalho, Anna Crist, Ania Pino-Querido, Boris Shraiman, Matthew V. Rockman, Henrique Teotónio

## Abstract

Understanding the genetic basis of complex traits remains a major challenge in biology. Polygenicity, phenotypic plasticity and epistasis contribute to phenotypic variance in ways that are rarely clear. This uncertainty is problematic for estimating heritability, for predicting individual phenotypes from genomic data, and for parameterizing models of phenotypic evolution. Here we report a recombinant inbred line (RIL) quantitative trait locus (QTL) mapping panel for the hermaphroditic nematode *Caenorhabditis elegans*, the *C. elegans* multiparental experimental evolution (CeMEE) panel. The CeMEE panel, comprising 507 RILs, was created by hybridization of 16 wild isolates, experimental evolution at moderate population sizes and predominant outcrossing for 140-190 generations, and inbreeding by selfing for 13-16 generations. The panel contains 22% of single nucleotide polymorphisms known to segregate in natural populations, and complements existing mapping resources for *C. elegans* by providing high nucleotide diversity across >95% of the genome. We apply it to study the genetic basis of two fitness components, fertility and hermaphrodite body size at time of reproduction, with high broad sense heritability in the CeMEE. While simulations show we should detect common alleles with additive effects as small as 5%, at gene-level resolution, the genetic architectures of these traits does not feature such alleles. We instead find that a significant fraction of trait variance, particularly for fertility, can be explained by sign epistasis with weak main effects. In congruence, phenotype prediction, while generally poor (*r*^2^ < 10%), requires modeling epistasis for optimal accuracy, with most variance attributed to the highly recombinant, rapidly evolving chromosome arms.

## Introduction

Most measurable features of organisms vary among individuals. Outlining the genetic dimension of this variation, and how this varies across populations and traits, has important implications for the application of genomic data to predict disease risk and agricultural production, for estimation of heritability, and for understanding evolution (Lynch and Walsh 1998; Barton and Keightley 2002). Complex traits are defined by being multifactorial. They tend to be influenced by many genes and to be plastic in the presence of environmental variation, and the manner in which phenotypic variation emerges from the combined effects of causal alleles is rarely clear. Although phenotype prediction and some aspects of evolution can often be well approximated by considering additive effects alone, non-additive interactions between alleles at different loci (with marginal additive effects) may explain a large fraction of trait variation yet remain undetected due to low statistical power (Phillips 2008). Adding further complication, one cannot usually assume that genetic and environmental effects are homogeneous or independent of one another (Barton and Turelli 1991; Félix and Barkoulas 2015), nor that the genetic markers used for mapping quantitative trait loci (QTL) are faithfully and uniformly associated with causal alleles (Yang *et al*. 2010; Speed *et al*. 2012).

Human height, for example, is the canonical quantitative trait, an easily measured, stable attribute with high heritability (around 80%) when measured in families Fisher (1930); Galton (1886); Visscher *et al*. (2010). Hundreds of common QTL (minor allele frequency, MAF>5%) of small effect have been detected by genome-wide association studies (GWAS) over the last two decades, explaining in sum only a small fraction (around 20%) of heritability (Wood et al. 2014). A recent study with more than 7 × 10^5^ people showed that close to one hundred uncommon QTLs (0.1%<MAF<5%) of more moderate effects explain a mere extra 5% of heritability (Marouli et al. 2017). It has taken methods of genomic selection in animal breeding, and dense genetic marker information (Meuwissen et *al*. 2001; Meuwissen and Goddard 2010), to show that common QTL of very small effect can potentially explain a large fraction of the variability in human height and common diseases (Yang *et al*. 2010;*Speed et al*. 2016). Thus, in perhaps many cases, the so-called problem of the “missing heritability” may be synonymous with high polygenicity (Hill et *al*. 2008; Manolio et *al*. 2009). The contribution of statistical epistasis to variation in human height is likely to be modest (Visscher et *al*. 2010), although the generality of this for size-related traits in other organisms is not known. Molecular genetics and biochemistry suggest functional non-additivity is ubiquitous within individuals, and significant effects on trait variation have been shown in many cases (e.g., MUKAI (1967); Whitlock and Bourguet (2000); Bonhoeffer *et al*. (2004); Carlborg *et al*. (2006); de Visser *et al*. (2009); Zwarts *et al*. (2011);*Shao et al*. (2008); Gaertner *et al*. (2012); Barkoulas *et al*. (2013); Weinreich *et al. (2013)*; Huang *et al*. (2014); Vanhaeren *et al*. (2014); Bloom *et al*. (2015); Monnahan and Kelly (2015b,a); Paaby *et al*. (2015); Tyler *et al*. (2016); Schoustra *et al*. (2016); Forsberg *et al*. (2017); Chirgwin *et al*. (2016), but the importance of epistasis in shaping fitness landscapes and in generating the additive genetic variance on which selection can act is still debated (Cheverud and Routman 1995; Wolf *et al*. 2000; Phillips 2008; Hansen 2013; Mackay *et al*. 2014)).

Alongside GWAS, inbred line crosses in model systems continue to be instrumental for our understanding of the genetics of complex traits, given the opportunity for control of confounding environmental covariates and accurate measurement of breeding values. Crosses among multiple parental strains in particular - such as those now available for mice (Churchill *et al*. 2004), Drosophila (Macdonald and Long 2007), maize *(McMullen et al*. 2009; Buckler *et al*. 2009), wheat (Huang *et al*. 2012;*Mackay et al*. 2014; Thepot *et al*. 2015), rice (Bandillo *et al*. 2013), tomato (Pascual *et al*. 2015) and Arabidopsis (Kover *et al*. 2009), among others - have been developed to better sample natural genetic variation. Greater variation also allows the effects of multiallelic loci to be studied and, subject to effective recombination, improved QTL resolution. If large populations and random mating are imposed for long periods, gains in resolution can be dramatic (Valdar *et al*. 2006; Rockman and Kruglyak 2008), although this comes at the expense of increased opportunity for selection to purge diversity (e.g., Baldwin-Brown *et al*. (2014); Rockman and Kruglyak (2009)).

Better known as a model for functional biology *(Corsi et al*. 2015), the nematode *Caenorhabditis elegans* has also contributed to our understanding of complex traits and their evolution. *C. elegans* shows extensive variation in complex traits (Gems and Riddle 2000; Knight *et al*. 2001; Barrière and Félix 2005; Gutteling *et al. 2007;* Gray and Cutter 2014; Diaz and Viney 2014; Teotónio *et al*. 2017) and sex-determination and breeding mode (selfing and outcrossing) can be genetically manipulated at will. QTL for traits such as embryonic lethality (Rockman and Kruglyak 2009), pesticide resistance (Ghosh *et al*. 2012) and telomere length (Cook *et al*. 2016) have been found by association studies in an ever expanding panel of inbred wild isolates, the *C. elegans* natural diversity resource (CeNDR; https://elegansvariation.org/, Cook *et al*. (2017)). QTL for a range of complex traits have also been found using collections of recombinant inbred lines (RILs) (Rockman and Kruglyak 2009) and introgression lines (ILs) (Doroszuk *et al*. 2009) derived from crossing the laboratory domesticated N2 strain (Sterken *et al*. 2015) and the divergent Hawaiian wild isolate CB4856 (e.g., Andersen *et al*. (2014, 2015)), or by two-parent crossing of non-domesticated strains (e.g., Duveau and Félix (2012); Noble *et al*. (2015)). GWAS and two-parent crosses have given insights into how natural selection has shaped phenotypic variation in *C. elegans* and related nematodes. For example, an N2/CB4856 RIL panel has been used to argue that selection on linked sites largely explains the distribution of QTL effects for mRNA abundance (Rockman *et al*. 2010). Lastly, *C. elegans* is also one of the main models for experimental evolution (Gray and Cutter 2014; Teotónio *et al*. 2017). Mutation accumulation line panels in particular have long been used to estimate mutational heritability (Estes and Lynch 2003; Estes 2005;*Baer et al*. 2005; Baer 2008; Phillips *et al*. 2009; Halligan and Keightley 2009) and to argue that standing levels of genetic variation in natural populations for complex traits can be explained by a mutation-selection balance (Etienne *et al*. 2015; Farhadifar *et al*. 2016). As yet, the QTL mapping resolution of existing *C. elegans* RIL panels has been coarse, and there is no panel derived from crosses of multiple wild parental strains.

A prominent characteristic of *C. elegans* is its mixed androdioecious reproductive system, with hermaphrodites capable of either selfing, from a cache of sperm produced late in larval development (Hirsh *et al*. 1976), or outcrossing with males (Maupas 1900). Sex determination is chromosomal, with hermaphrodites XX, and XO males maintained through crosses and rare X-chromosome non-disjunction during hermaphrodite gametogenesis (Nigon 1949). Because males are typically absent from selfed broods but are half the progeny of a cross, twice the male frequency in a population is the expected outcrossing rate (Stewart and Phillips 2002; Cutter 2004). Natural populations have low genetic diversity and very high linkage disequilibrium (LD), with generally weak global population structure and high local diversity among typically homozygous individuals at the patch scale (Barrière and Félix 2005, 2007; Cutter *et al*. 2009). Average single nucleotide polymorphism (SNP) diversity is on the order of 0.3% (Cutter 2006) though highly variable across the genome, reaching 16% or more in some hypervariable regions (Thompson *et al*. 2015). Low diversity and high LD is due to the predominance of inbreeding by selfing, which reduces the effective recombination rate and elevates susceptibility to linked selection (Rockman *et al*. 2010; Andersen *et al*. 2012). Crosses between wild isolates have revealed outbreeding depression (Dolgin *et al*. 2007; Chelo *et al*. 2014), which may be in part due to the disruption of epistatic allelic interactions. Evidence supporting this prediction in *C. elegans* is, to date, scarce: one study has shown that recombination between several QTL “complexes” leads to dysregulation of thermal preferences *(Gaertner et al*. 2012).

Although selfing is the most common reproductive mode in natural *C. elegans* populations, males, though rare, are variably proficient in mating with hermaphrodites (Teotónio *et al*. 2006; Murray *et al*. 2011). Perhaps as a consequence of low but significant outcrossing (and also a metapopulation demographic structure) several loci have been found to be under some form of balancing selection (e.g., Ghosh *et al*. (2012); Greene *et al*. (2016)). Moreover, evolution experiments involving crosses among multiple strains have shown that high outcrossing rates can persist as long as there is heritable variation for male traits (Anderson *et al*. 2010; Teotónio *et al*. 2012; Masri *et al*. 2013). In our evolution experiments in particular (Teotónio *et al*. 2012), moderate population sizes and high outcrossing rates facilitated the loss of genetic diversity by (partial) selective sweeps, with excess heterozygosity maintained by epistatic selection on overdominant loci (e.g., Chelo and Teotónio (2013); Chelo *et al*. (2014)).

This foundation suggests study of *C. elegans* may be fruitful for our understanding of the contribution of within- and between-locus interactions to complex traits and their evolution. Here we present a panel of 507 genome sequenced RILs obtained by intercrossing 16 wild isolates, culturing at high out-crossing rates in populations of ≈ 10^4^ for 140-190 generations of experimental evolution, followed by inbreeding by selfing for 13-16 generations. The *<u>C.</u> elegans* <u>M</u>ultiparental <u>E</u>xperimental <u>E</u>volution (CeMEE) RIL panel complements existing *C. elegans* mapping resources by providing fine mapping resolution and high nucleotide diversity. Using simulations, we show that the CeMEE panel can give gene-level resolution for common QTL with effects as low as 5%. In subsets of the CeMEE, we investigate the genetic basis of two fitness components, fertility and hermaphrodite body size at the time of reproduction, by variance decomposition under additive and additive-by-additive epistatic models, and by genome-wide 1- and 2-dimensional association testing. We find that the genetic basis of both traits, particularly fertility, is highly polygenic, with a significant role for epistasis.

## Materials and Methods

### CeMEE derivation

The panel was derived in 3 stages (Figure 1). First, 16 wild isolates (AB1, CB4507, CB4858, CB4855, CB4852, CB4586, MY1, MY16, JU319, JU345, JU400, N2 (ancestral), PB306, PX174, PX179, RC301; obtained from the Caenorhaditis Genetics Center) were inbred by selfing for 10 generations to ensure homozygosity, then intercrossed to funnel variation into a single multiparental hybrid population, as described in Teotónio *et al*. (2012). Each of the four funnel phases comprised multiple pairwise, reciprocal crosses at moderate population sizes (see Figure S1 of Teotónio *et al*. (2012) for full details of replication and population sizes).

**Figure 1.**
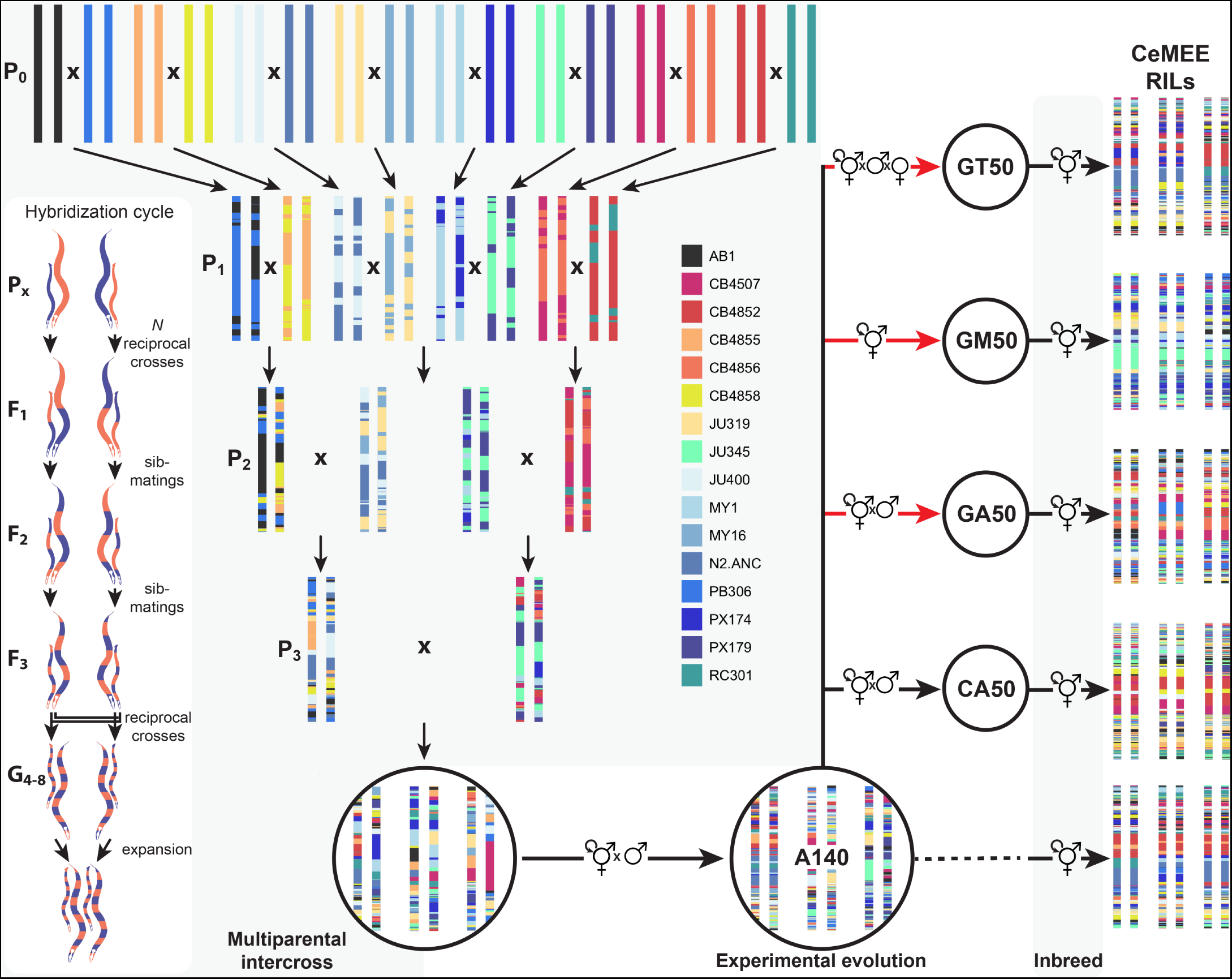
CeMEE derivation. The multiparental intercross funnel phase comprised four stages of pairwise crosses and progeny mixing, carried out in parallel at controlled population sizes. One hybridization cycle for a single founder cross is inset at left: in each cycle, multiple reciprocal crosses were initiated, increasing in replicate number and census size each filial generation. *F*_1_ and *F*_2_ progeny were first sib-mated, then reciprocal lines were merged by intercrossing the F3 and expanding the pooled *G*_4_ (for three to four generations) before commencing the next reduction cycle. The resulting multiparental hybrid population was archived by freezing, and samples were thawed and then maintained for 140 non-overlapping generations of mixed selfing and outcrossing under standard laboratory conditions to generate the A140 population. Hermaphrodites were then sampled from the A140 and selfed to generated the A140 RILs. Additionally, the outbred A140 population was evolved for a further 50 generations under the same conditions (control adapted lines; CA) or under adaptation to a salt gradient with varying sex ratios (GT, GM and GA lines; Theologidis *et al*. (2014)). See Materials and Methods for description of sub-panels, and Teotónio *et al*. (2012) for details of replicate numbers and population sizes for each funnel generation.

Second, the multiparental hybrid population was evolved for 140 discrete generations at population sizes of *N* ≈ 10^4^ (out-crossing rate ≈ 0.5, *N*_*e*_ ≈ 10^3^), to obtain the A140 population, as reported in (Teotónio *et al*. 2012; Chelo and Teotónio 2013; Chelo *et al*. 2013). Sex-determination mutations were then mass introgressed into the A140, while maintaining genetic diversity, to generate monoecious (obligately selfing hermaphrodites) and trioecious (partial selfing with males, females and hermaphrodites) populations, as detailed in Theologidis *et al*. (2014). Further replicated experimental evolution was carried out for 50 generations under two environmental regimes: (1) a <u>C</u>ontrol regime (conditions as before), with the wild-type <u>A</u>ndrodioecious reproductive system (CA50 collectively, full designations can be found in Table S1); and (2) a <u>G</u>radual exposure to an increasing gradient of NaCl, from 25mM (standard NGM-lite medium, US Biological) to 305mM until generation 35 and thereafter, varying reproductive system (GX50, where *X* is <u>A</u>ndrodioecious, <u>M</u>onoecious or <u>T</u>rioecious). Although trioecious populations started evolution with only 0.1% of hermaphrodites, by generation 50 they were abundant (50%; see Figure S7 in Theologidis *et al*. (2014)). Androdioecious populations maintained outcrossing rates of >0.4 until generation 35, soon after losing males to finish with an outcrossing rate of about 0.2 by generation 50 (Figure S5 in Theologidis *et al*. (2014)). The effects of reproductive system on the genetics and evolution of complex traits will be the subject of future work.

Finally, hermaphrodites were inbred by selfing to obtain recombinant inbred lines (RILs). Population samples (> 10^3^ individuals) were thawed from -80C and maintained under standard laboratory conditions for two generations. At the third generation, single hermaphrodites were picked at the late third to early fourth (L3/L4) larval stage and placed in wells of 12-well culture plates, containing M9 medium (25mM NaCl) seeded with *E. coli*. Lines were propagated at 20C and 80% RH by transferring a single L3/L4 individual for 16 (A140 population) or 13 generations (4-7 days between transfers). At each passage, parental plates were kept at 4C to prevent growth until offspring production was verified, and in the case of failure a second transfer was attempted before declaring line extinction. Inbreeding was done in several blocks from 2012 to 2016, in two different labs. A total of 709 RILs were obtained and archived at -80C (File S2).

### Sequencing and genotyping

DNA of the 16 founders, 666 RILs and the A140 population was prepared using the Qiagen Blood and Tissue kit soon after derivation or after thawing from frozen stocks and expansion to at least 10^4^ L1 individuals. Founders were sequenced to >=30X depth with 50 or 100bp paired-end reads (Illumina HiSeq 2000, New York University Center for Genomics and Systems Biology GenCore facility). Reads were mapped (BWA 0.7.8; Li and Durbin (2010)) to the WS220 *C. elegans* N2 reference genome and variants (SNPs and small indels) were called jointly (GATK 3.3-0 HaplotypeCaller; McKenna *et al*. (2010)), followed by base quality score recalibration (BQSR) using a subset of high scoring sites (29% of initial variants passing strict variant filtration: “MQ < 58.0 | | DP <20 | | FS > 40.0 | | SOR > 3.0 | | ReadPosRankSum < -5.0 | | QD < 20.0 | | DP > mean × 2”). Diallellic single nucleotide variants on the six nuclear chromosomes were intersected with calls from a joint three-sample call (GATK UnifiedGenotyper) on pooled founders, a subset of pooled RILs (SUP TABLE XX, SAME AS CEMEE LIST ANOTHER COLUMN), and 72X sequencing of the A140 population (approximately 1400x total), then filtered based on variant call metrics (MQ < 50.0 | | DP < 10 | | FS > 50.0 | | SOR > 5.0 | | ReadPosRankSum < -5.0 | | QD < 6.0 | | DP > mean × 3) and on the number of heterozygous or missing founder calls (3,014 sites > 8 removed; these calls likely represent copy number differences between founders and the N2 reference), and requiring ≥ 1 homozygote (28,872 sites removed), giving an initial set of 404,536 SNP markers.

RILs were sequenced with 100bp paired-end reads (Nextera libraries, HiSeq 2000, NYU) or 150bp paired-end reads (Hiseq X Ten, BGI Tech Solutions Company, Hong Kong), to mean depth 7.2X (minimum 0.2X). Genotypes were imputed by Hidden Markov Model (HMM) considering the 16 founder states and mean base qualities of reads. Downsampled predictions for a subset of RILs sequenced to high (20-30X) depth gave imputation accuracy of approximately 99% at 0.2X and 99.9% at 0.5X (93% of lines).

We assessed accuracy and appropriate variant filtering thresholds by genotyping a set of 784 markers, uniformly distributed across the six chromosomes according to the genetic distances of Rockman and Kruglyak (2009), in 182 RILs with the iPlex Sequenom MALDI-TOF platform (Bradic *et al*. 2011). Sequenom data can be found in Table S2. We fitted a linear model with counts of Illumina/Sequenom concordant cases as the response variable, and all founder variant quality metrics together with the number of missing or heterozygous calls in the founders, the number of zero-coverage or potentially heterozygous sites (with at least a single Illumina read for each genotype), variant nucleotide identity, and reference nucleotide and dinucleotide identity as explanatory variables. Concordance across sequencing platforms was 96.9% after (93.7% before) final filtering, and we retained 388,201 diallelic SNPs as founder markers. We estimated residual heterozygosity for 25 A140 lines sequenced to >20X coverage (single sample calls using GATK 3.3-0 HaplotypeCaller, variant filtration settings MQ < 50.0 | | DP < 5 | | MQRankSum < -12.5 | | SOR > 6 | | FS > 60.0 | | ReadPosRankSum < -8.0 | | QD < 10.0 | | DP > mean×3). Mean heterozygosity at founder sites is 0.095% (standard deviation 0.042%, range 0.033-0.18%).

After removal of RILs sharing greater than the mean pairwise identity + 5 standard deviations (84.8%, excluding monoecious lines), we retained 178 A140 RILs, 118 CA50 RILs (from three replicate populations), 127 GA50 RILs (three replicates), and 79 GT50 RILs (two replicates). The 98 GM50 RILs (two replicates) are highly related on average and group together into a small number of “isotypes”. To prevent introduction of strong structure, we discard all but five below the above panel-wide pairwise identity threshold for the purposes of trait mapping. In total, the CeMEE comprises 507 RILs from five sub-panels, with 352,583 of the founder markers segregating within it (File S3).

### CeMEE genetic structure

#### Differentiation from natural isolates and founders

We compared similarity within and between the CeMEE RILs and 152 sequenced wild-isolates from the CeNDR panel (release 20160408). The distributions for all pairwise genotype and haplotype (% identity at 0.33cM scale in F2 map distance) distances are plotted in Figure 2, for 256,535 shared diallelic sites with no missing or heterozygous calls.

**Figure 2.**
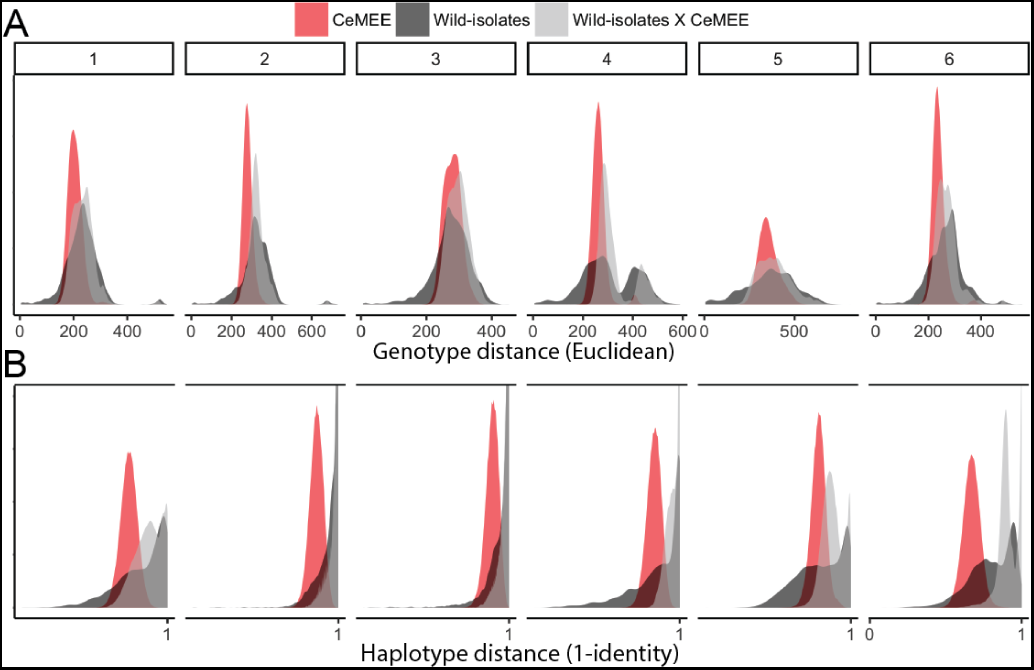
Similarity among CeMEE RILs and 152 sequenced wild-isolates (*Caenorhabditis elegans* Natural Diversity Resource) at 256,535 shared diallelic sites. The distribution of pairwise genotype (**A**) and haplotype (**B**) distances, within and between CeMEE RILs and CeNDR wild-isolates, by chromosome. Haplotype distances are 1-% identity at 0.1cM scale. Note that chromosomes 2-4 all show a marked excess in haplotype dissimilarity between CeMEE RILs and CeNDR wild-isolates, and the density is truncated by a factor of four for visibility.

Linkage disequilibrium (*r*^2^) was computed for founders and CeMEE RILs at the same set of sites (MAF >1/16, <5% ambiguous imputed RIL genotypes and ≤ 1 heterozygous/missing founder genotypes, then downsampled by 10 for computational tractability), and plotted against genetic distances (obtained by linear interpolation from the N2/CB4856 map, scaled to *F*_2_ distances (Rockman and Kruglyak 2009). To assess the extent of subtle, long-range linkage disequilibrium in the form of inter-chromosomal structure, we compared *r*^2^ among chromosomes to a null distribution generated by permutation (*n*=5000). In each permutation, filtered RIL genotypes (pruned of strong local linkage *r*^2^ < 0.98, no ambiguous calls) were randomly down-sampled to equal size across chromosomes, split by chromosome, then shuffled within each sub-panel before taking the mean correlation across chromosomes (or omitting all single and pairwise chromosome combinations) as test statistic. The effect of local LD pruning is to reduce the weighting of large regions in strong linkage in order to better assay weak interactions across the remainder of the genome.

#### Reconstruction of ancestral haplotypes and genetic map expansion

For each RIL, founder haplotypes were inferred with the RABBIT HMM framework implemented in Mathematica (Zheng et al. 2015), conditioning on the recombination frequencies observed for the N2 × CB4856 RILs (scaled to *F*_2_ map length) (Rockman and Kruglyak 2009). Realized map expansion was estimated by maximum likelihood for each chromosome, before full marginal reconstruction of each chromosome (explicitly modeling recombination on the *X* and autosomes) using posterior decoding under the fully dependent homolog model (dep-Model). Under this model, appropriate for fully inbred diploids, chromosome homologs are assumed to have identical ancestral origins (prior identity by descent probability *f* = 1), and the recombination junction density (transition probability) is given by the estimated map expansion (*Ra*) and genotyping error rates (set to 5×10^−5^ for founders and 5×10^−3^ for RILs based on likelihood from a parameter sweep). Sites called as heterozygous or missing in the founders, or unresolved to [0, 1] by the genotype imputation HMM were set to NA before reconstruction. For reconstruction summaries, haplotype posterior probabilities were filtered to >0.2, and haplotype lengths and breakpoints were estimated from run lengths of marker assignments, taking the single best haplotype (if present), maintaining haplotype identity (if multiple assignments of equal probability), or the first among equals otherwise.

To test reconstruction accuracy as a function of haplotype length, we performed simulations of a pedigree varying only the number of generations of random mating. Starting from a single population representing all founders (*N*=1000, corresponding to the expected *N_e_* during experimental evolution), mating occurred at random with equal contribution to the next generation. Recombination between homologous chromosomes occurred at a rate of 50cM, with full crossover interference, and the probability of meiotic crossover based on distances between marker pairs obtained by linear interpolation of genetic positions (Rockman and Kruglyak 2009). For each chromosome, 10 simulations were run sampling at 10, 25, 50, 100 and 150 generations, and haplotype reconstruction was carried out as above. Maximum likelihood estimates of realized map expansion for simulations were used to calibrate a model for prediction of realized number of generations in the RILs by chromosome. A 2nd degree polynomial regression of *Ra* on the known number of generations was significantly preferred over a linear fit by likelihood ratio test, given significant underestimation as pedigree length increased (approaching 10% at *G*_150_).

#### Population stratification

Population stratification was assessed using (1) principal component decomposition, giving a uni- or bivariate view of the importance of genetic structure associated with CeMEE sub-panels, and (2) by supervised and unsupervised discriminant analysis of principal components (DAPC; Jombart et al. (2010)), giving an estimate of the fraction of principal component variance that best predicts sub-panel structure, and an inference of population structure without regard to sub-panel identities. In all cases decomposition was of scaled and centered genotypes pruned of strong local LD (*r*^2^ < 0.98), giving all markers equal weight (and therefore more weight to low frequency alleles).

Of the first 50 principal components, 10 are significantly associated with sub-panel identity (i.e., evolutionary history) by ANOVA (*p* < 0.05 after Bonferroni correction), accounting for just 3.9% of the variance in sum. Seven of the top 10 PCs are significant, though others up to PC 38 are also associated, showing that multiple sources of structure contribute to the major axes of variation. Fitting all pairs among the the top 50, two pairs (7 and 19, 13 and 14) are significant (again at a conservative Bonferroni adjusted threshold), resolving the GT50 RILs as most distinct.

For DAPC (R package adegenet, Jombart (2008)), we used 100 rounds of cross-validation to determine the number of principal components required to achieve optimal group assignment accuracy (the mean of per-group correct assignments). This value (40 PCs) was then used to infer groups by unsupervised *k*-means clustering (default settings of 10 starts, 10^5^ iterations), with k selected on the Bayesian Information Criterion (BIC). Correspondence of inferred groups with known groups was tested by permutation. Given the contingency table C, where *C*_*i,j*_ represents the number of lines known to be in sub-panel *i* and inferred to be in cluster *j*, the inferred values for each cluster (*js*) were shuffled among known groups (*is*) 10,000 times, with the sum of the variance among known groups taken as a summary statistic (high values reflecting significant overlap between inferred and known groups).

### Phenotyping

#### Fertility

In the experimental evolution scheme under which the CeMEE RILs were generated, a hermaphrodite’s contribution to the next generation is the number of viable embryos that survive bleaching (laid, but unhatched, or held *in utero*) that subsequently hatch to L1 larvae 24h later. We treat this phenotype as fertility, and measured it for individual worms of 230 RILs. Each line was thawed and maintained for two generations under standard conditions (Stiernagle 2006; Teotónio *et al*. 2012; Theologidis *et al*. 2014), bleached to kill adults, then embryos were allowed to hatch and synchronize as L1 larvae. L1s were then moved to fresh plates seeded with *E. coli* and allowed to develop for 48 hours. Single L3-L4 staged hermaphrodite larvae were then placed into each well of 96-well plates using a micropipette and stereomicroscope. Plate wells contained NGM-lite + 100μg/ml ampicillin, previously inoculated with 1*μ*l of an overnight culture of *E. coli* (HT115) and stored until usage at 4C (maximum 2 weeks before use). After transfer, plates were covered with Parafilm to prevent cross-contamination and incubated at 20C and 80% relative humidity (RH) until the following day. Embryos were extracted by adding bleach solution to wells (1M KOH, 5% NaClO 1:1 v/v in M9 buffer) for 5 minutes, then 200*μ*l of the extract was removed and rinsed 3 times in M9 buffer by centrifugation. The M9 suspension (200*μ*l) was then transferred to another 96-well plate containing 120*μ*l of M9 per well. Plates were incubated overnight (as above), then centrifuged for 1 min at 1800rpm to sediment any swimming larvae before imaging at 4 pixel/*μ*m^2^ with a Nikon Eclipse TE2000-S inverted microscope. ImageJ was then used to manually count the number of live (moving) L1s in each well. During assay setup and image analysis wells were censored where: bacteria were absent; hermaphrodites were absent or dead at the time of bleach; males had been inadvertently picked; more than 1 adult was present; or hermaphrodites had not been killed upon bleaching. Except for density between the L4 stage until reproduction, all assay conditions were the same as those used during experimental evolution. Fertility measurements do not include potential survival differences between the L1 stage until reproduction, but we nonetheless take it as a surrogate for fitness (Chelo *et al*. 2013).

Two independent plates within a single thaw were set-up for most RILs (1 plate for six lines, maximum=4, mean=2.0), which we consider as replicates for estimation of repeatability (see below). In total the median number of measurements per line was 43 (range 4-84). Highly replicated data for the reference strain N2 were also included for modeling purposes (404 observations across 17 plates, spanning 9 of 47 independent thaws). Wells with no offspring were observed for 4% of N2 data (and 2.9% of all RIL data). These are likely to be due to technical artifact, such as injury or incorrect staging, and were excluded before modeling. Mapping values were the Box-Cox transformed line coefficients from a Poisson generalized linear model with fixed effects of plate row, column and edge (exterior rows and columns), and the count of offspring per worm as response variable. Three outliers with coefficients >3 standard deviations below the mean were excluded, leaving data for 227 RILs (File S4). Data come from RILs of three sub-panels (170 A6140, 45 GA50, 12 GT50), which explains 4% of trait variation (GA50 RILs have higher mean fertility than the A6140, regression coefficient = 0.43, *p* = 0.01; see Figure S1).

#### Adult hermaphrodite body size

412 RILs were thawed and maintained for two generations under standard conditions. On the third generation, 1000 synchronized L1 larvae were moved to NGM-lite plates (25mM NaCl) where they developed and matured for 3 days. Image data was acquired at the usual time of reproduction (as during experimental evolution) and analysed with the Multi-Worm Tracker (Swierczek et al. 2011), using a Dalsa Falcon 4M30 CCD camera and Schott backlight A08926. Tracking was performed for 25 minutes with default parameters, and particle (worm) contours extracted (on average, 300 particles obtained every 0.5s). Raw values from each plate were calculated from track segments of length 40-41s taken at 80s intervals, ultimately estimating the area of an individual as the grand mean of the per-segment estimates (accounting for temporal autocorrelation within a time-series, analysis not shown).

Assays were carried out in two lab locations over several years, while recording the relative humidity and temperature at the time of assay. Mapping values are the Box-Cox transformed line coefficients from a linear model incorporating fixed effects of year, nested within location, and humidity and temperature, nested within location. Data come from a mean of 2.1 (maximum 4) independent thaw blocks for each RIL, for 410 RILs after excluding 2 outliers >3 standard deviations below the mean, with a median of 447 measurements per RIL and block (range 109-1013; File S5). Data for the reference strain N2 were also included in the model (1664 observations from two plates). Data come from RILs of three sub-panels (165 A6140,118 CA50,127 GA50), which explains 17% of trait variation (GA50 RILs are much larger than the A6140, regression coefficient = 0.94, p < 10^−16^; see Figure S1). This difference is not obviously associated with technical covariates, since data acquisition for A140 RILs and GA50 RILs was distributed similarly with respect to location and time.

Fertility and body size are moderately correlated (Figure S1; see also Poullet et al. (2016)), justifying the latter being considered a fitness-proximal trait (Spearman’s *ρ* = 0.354, *p* = 2.336 × 10^−7^ for mapping coefficients, for 202 lines with data for both traits).

### Heritability and phenotype prediction

#### Repeatability

Repeatability was estimated from ANOVA of the line replicate means for each trait as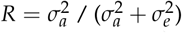, where 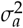= (mean square among lines -mean square error)/*n*_0_, and *n*_0_ is a coefficient correcting for varying number of observations (1-4 plate means) per line (Lessells and Boag 1987; Sokal and Rohlf 1995). Assuming equal variance and equal proportions of environmental and genetic variance among replicates, *R* represents on upper bound on broad-sense heritability (Falconer 1981; Hayes and Jenkins 1997). Fertility data were square root transformed to decouple the mean and variance.

#### Assumptions

In inbred, isogenic, lines, broad-sense heritability can also be estimated by linear mixed effect model from the covariance between genetic and phenotypic variances. The measurement of genetic similarity is, however, subject to a number of assumptions and is (almost) always, at best, an approximation (Speed and Balding 2015).

A first assumption is that all markers are the causal alleles of phenotypic variation. It is unavoidable, however, that markers tag the (unknown) causal alleles to different degrees due to variable linkage disequilibrium. A second, usually implicit, assumption in calculating genetic similarity is the weight given to markers as a function of allele frequency. Equal marker weights have commonly been used in animal breeding research, while greater weight has typically been given to rare alleles in human research, which has some support under scenarios of both selection and neutrality (Pritchard 2002). A third assumption, related to the first two, is the relationship between LD and causal variation. If the relationship is positive -causal variants being enriched in regions of high LD - then heritability estimated from all markers will be upwardly biased, since the signal from causal variation contributes disproportionately to genetic similarity (Speed *et al*. 2012).

The use of whole genome sequencing largely addresses the first assumption, given (as here) very high marker density and an accurate reference genome, although in the absence of full *de novo* genomes from long-read data for each individual, the contribution of large scale copy-number and structural variation, and new mutation, will remain obscure. To account for the second and third assumptions, we used LDAK (v5.0) to explicitly account for LD in the CeMEE (decay half-life = 200Kb, min-cor = 0.005, min-obs = 0.95) (Speed *et al*. 2012). Heritability estimates were not sensitive to variation in the decay parameter over a 10-fold range or to the measurement unit (physical or genetic), although model likelihoods were non-significantly better for physical distance. Across the set of 507 RILs, 88,508 segregating markers were used after local LD-based pruning (*r*^2^<0.98) and, of these, 22,984 markers received non-zero weights. LD-weighting can magnify the effects of genotyping errors. We excluded 17,740 markers with particularly low local LD (mean *r*^2^ over a 20 marker window < 0.3, or the ratio of mean *r*^2^ to that of the window mean < 0.3). Heritability estimates were largely unchanged (within the reported intervals), as were our general conclusions on variance components and model performance.

#### Modeling

Model fit was assessed by phenotype predictions from leave-one-out cross validation, calculating the genomic best linear unbiased prediction (GBLUP; Meuwissen *et al*. (2001); VanRaden (2008); Yang *et al*. (2010)) for each RIL and returning the squared correlation coefficient (*r*^2^) between observed and predicted trait values. To avoid bias associated with sample size all models were unconstrained (non-error variance components were allowed to vary outside 0-1 during convergence) unless otherwise noted, which generally gave better fit for multi-component models.

Given *m* SNPs, genetic similarity is calculated by first scaling S, the n × m matrix of mean centered genotypes, where *S_i,j_* is the number of minor alleles carried by line *i* at marker *j* and frequency *f*, to give *X*: 
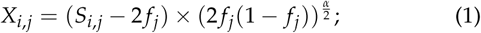

The additive genomic similarity matrix (GSM) **A** is then ***XX*^*T*^** / m. Here *α* scales the relationship between allele frequency and effect size (Speed *et al*. 2012). *α* = −1 corresponds to the assumption of equal variance explained per marker (an inverse relationship of effect size and allele frequency), while common alleles are given greater weight at α>0. We tested α ∈ [−1.5, −1, −0.5,0,0.5,1] and report results that maximized prediction accuracy. With *Υ* the scaled and centered vector of n phenotype values, the additive model fit for estimating genomic heritability *h^2^* is then: 
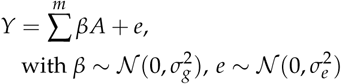
where *β* represents random SNP effects capturing genetic variance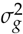, *e* is the residual error capturing environmental variance 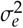. Given *Υ* and **A**, heritability can be estimated from restricted/residual maximum likelihood (REML) estimates of genetic and residual variance as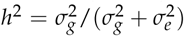. Note that we use the terms *h*^2^ and genomic heritability interchangeably here for convenience, although in some cases the former includes non-additive covariances. We assume RILs are fully inbred, and so dominance variance does not contribute to heritability.

The existence of near-discrete recombination rate domains across chromosomes has lead to characteristic biases in nucleotide variation, correlated with gene density and function (Cutter *et al*. 2009). Similarly, recent selective sweeps, coupled with the low effective outcrossing rate in *C. elegans*, have lead to a markedly unequal distribution of variation across chromosomes (Andersen *et al*. 2012; Rockman *et al*. 2010). This variability in mutational effect, along with variable LD in the RILs, is not captured by aggregate genome-wide similarity with equal marker weighting (Speed *et al*. 2012; Goddard *et al*. 2016). We therefore first tested genetic similarity by explicitly modeling observed LD (Speed *et al*. 2012), with markers weighted by the amount of genetic variation they tag along chromosomes, and by their allele frequency (see above). Given *m* weights reflecting the amount of linked genetic variation tagged by each marker, *w*_*i*_,…, *w*_*m*_, the variance covariances for the basic model become: 
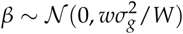
where W is a normalizing constant. Second, we jointly measured the variance explained by individual chromosomes (and by recombination rate domains within each chromosome), which can further improve the precision of heritability estimation if causal variants are not uniformly distributed by allowing variance to vary among partitions. Third, we tested epistatic as well as additive genetic similarity with (1) the entrywise (Hadamard) product of additive GSMs, giving the probability of allele pair sharing (Henderson 1985; Jiang and Reif 2015), (2) higher exponents up to fourth order interactions and (3) haplotype-based similarity at multi-gene scale. Additional similarity components (additive or otherwise) are added as random effects to the above model to obtain independent estimation of variance components:
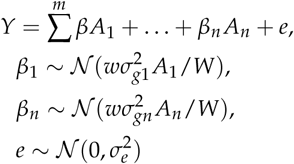

Haplotype similarity was calculated as the proportion of identical sites among lines at 0.033 and 0.067cM scales (corresponding to means of approximately 5 and 10Kb non-overlapping block sizes, or one and two genes), using either the diallelic markers only, or all called SNPs and indels. In the latter case, variants were imputed from reconstructed haplotypes if the most likely haplotypes of flanking markers were in agreement.

### GWAS

#### 1-dimensional tests

For single trait, single marker association, we fitted linear mixed models using the Python package LIMIX (https://github.com/PMBio/limix): 
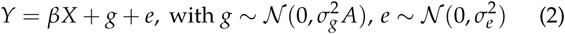
where *X* is the matrix of fixed effects (the SNP genotype of interest) and *β* is the effect on phenotypic variation that is estimated. *g* are the random effects describing genetic covariances (as above) accounting for non-independence among tests due to an assumed polygenic contribution to phenotype, with **A** the *n* × *n* genetic similarity matrix from the most predictive additive fit found for each trait above, and *e* is the error term.

To test the mapping resolution and power of the CeMEE panel, we carried out GWAS according to the model above for simulated phenotypes. We modeled a single focal additive locus (with *h^2^* from 1 to 30%) and a background polygenic component of equal variance (with scenarios of 10, 100 or 1000 loci), selected at random from SNPs with MAF > 0.05, and with genetic and environmental effect sizes drawn independently from the standard normal distribution. GWAS was carried out 1000 times for each scenario, controlling for relatedness with LD-weighted additive genetic similarity (α = -0.5). Power was estimated from a binomial generalized linear model considering all three polygenic scenarios together. Recall, the proportion of true positives passing significance, was assessed after masking a 1cM window around the focal SNP. 2-LOD drop intervals around the focal locus were calculated from similarly powered markers with ≥ MAF, with p-values converted to LOD scores as χ^2^/2 × *log*(2)/*log*_10_(2)).

For simulated traits all 507 lines and 262,218 markers (MAF > 0.05) were used for GWAS. For body size GWAS 410 lines and 254,174 markers were used, and 227 lines and 254,240 markers were used for fertility. Significance thresholds were established by permutation, with phenotypes generated by permuting phenotype residuals, given the estimated relatedness among lines (A), using the R package mvnpermute (Abney 2015). Significance level *α* is the corresponding percentile of the minimum p-values from 1000 permutations.

Given the correlation between traits (see above), we also tested a model for each trait on phenotype residuals after linear regression on the other, and a multi-trait model fitting effects common or specific to a trait. No markers passed significance in any case (analysis not shown).

#### 2-dimensional tests

We tested for additive-by-additive epistasis on the assumption of complete homozygosity. We first reduced the search space by local LD pruning (*r*^2^ < 0.5), requiring MAF > 0.05, the presence of all four two-locus homozygote classes at a frequency of ≥ 3, with ≤ 5 missing or ambiguous imputed genotypes (which were excluded from analysis). This gave a total of 19,913,422 tests for fertility (both inter- and intrachromosomal) and 28,138,090 for size, across 9,628 and 10,329 markers respectively. We tested for main and interaction additive effects for all marker pairs by ANOVA, taking as summary statistics the *F*-statistic for genotype interaction (2D tests), and also the sum of interaction scores for each marker (2D sum tests) above each of three thresholds (*F*>0,8,16, the latter corresponding roughly to the most significant single marker associations seen for both traits). All statistics were calculated separately for inter- and intrachromosomal tests. 2D sum tests are testing for excess weak to moderate interactions due to polygenic epistasis.

For computational tractability, tests were run in parallel on two chromosomes at a time. Null permutation thresholds were generated by shuffling phenotypes (using mvnpermute as above to ensure exchangeability in the presence of polygenicity or structure). 2D test thresholds were calculated for each chromosome separately from at least 2000 permutations each and differed little across chromosomes (α = 10%, 2.86 − 1.16 χ 10^−7^ for fertility, 1.86 × 10^−7^ - 7.2 × 10 ^−8^ for size). Inter- and intrachromosomal thresholds were calculated separately, but the reported interactions do not change if we pool both classes (or all chromosomes). 2D sum test thresholds were calculated separately for each chromosome pair and class (inter- and intrachromosomal).

We initially ignored relatedness for 2D testing, then fit linear mixed effect models as above with genetic covariance A for candidate interactions (R package hglm; Shen *et al*. (2014)). For size, the two candidate interactions all decreased slightly in significance (to a maximum p-value of 7.8 × 10^−7^), while significance increased for all four fertility interactions. The amount of phenotypic variance explained by candidates for each trait was estimated by ANOVA, jointly fitting all main and two-locus interactions.

### Data Availability

Sequence data are available from NCBI SRA under accession XXXXX. All data and methods scripts are archived in Dryad.org doi: XXX. RILs are available from the authors.

## Results and Discussion

### CeMEE differentiation from natural populations

The CeMEE panel of recombinant inbred lines draws variation from sixteen founders, and shuffles the diversity they contain through more than 150 generations at moderate population sizes and predominant outcrossing. The wild founders used to create the panel together carry approximately 25% of single nucleotide variants known to segregate in the global C. *elegans* population (CeNDR;*Caenorhabditis elegans* Natural Diversity Resource; Cook *et al*. (2017)). They vary, however, in distance to the N2 reference strain, with the Hawaiian CB4856 and German MY16 isolates together contributing over half of all markers, while the Californian CB4507 is closely related to N2 (Figure S3). Comparison of pairwise genetic distances in the CeMEE and 152 sequenced wild isolates (including a small number of more recently isolated, highly divergent lines) illustrates the extent of novelty generated by the multiparental cross (Figure 2). The CeMEE RILs occupy a substantial sub-space of the CeNDR genotypic diversity (Figure 2A), without the extensive haplotype sharing among wild-isolates and with the creation of many new multigenic haplotypes (Figure 2B).

### CeMEE differentiation from parental founders

Since *C. elegans* natural isolates suffer from outbreeding depression (?Gimond *et al*. 2013), the mixing phase is expected to generate high variance in fitness which, channeled through bottlenecks during serial intercrossing and population expansion, gives ample opportunity for loss of diversity through drift and selection. Fixation of N2 alleles at one X chromosome locus, spanning the known major effect behavioral locus *npr-1* (de Bono and Bargmann 1998; Gloria-Soria and Azevedo 2008; McGrath *et al*. 2009; Reddy *et al*. 2009; Andersen *et al*. 2014; *Bendesky et al*. 2011), during establishment of the A140 population has been documented with a coarse marker set (Teotónio *et al*. 2012). More broadly, the outbred A140 population showed non-negligible departure from the founders, with 32,244 alleles lost (unseen in both the A140 and RILs, 26,593 of these being founder singletons; Figure 3). Subsequent change during the inbreeding (and further adaptation) stages to generate RILs was more restricted, with an additional 3,171 alleles lost (2,542 of these at <10% frequency in both founders and the A140). Importantly, however, the physical distribution of allelic loss is relatively restricted: at least one marker is segregating in the CeMEE RILs at >5% minor allele frequency within 95.5% of 20Kb segments across the genome (97.2% of autosomal segments; for reference, protein coding genes are spaced just under 5Kb apart on average in the 100Mb *C. elegans* N2 genome).

**Figure 3.**
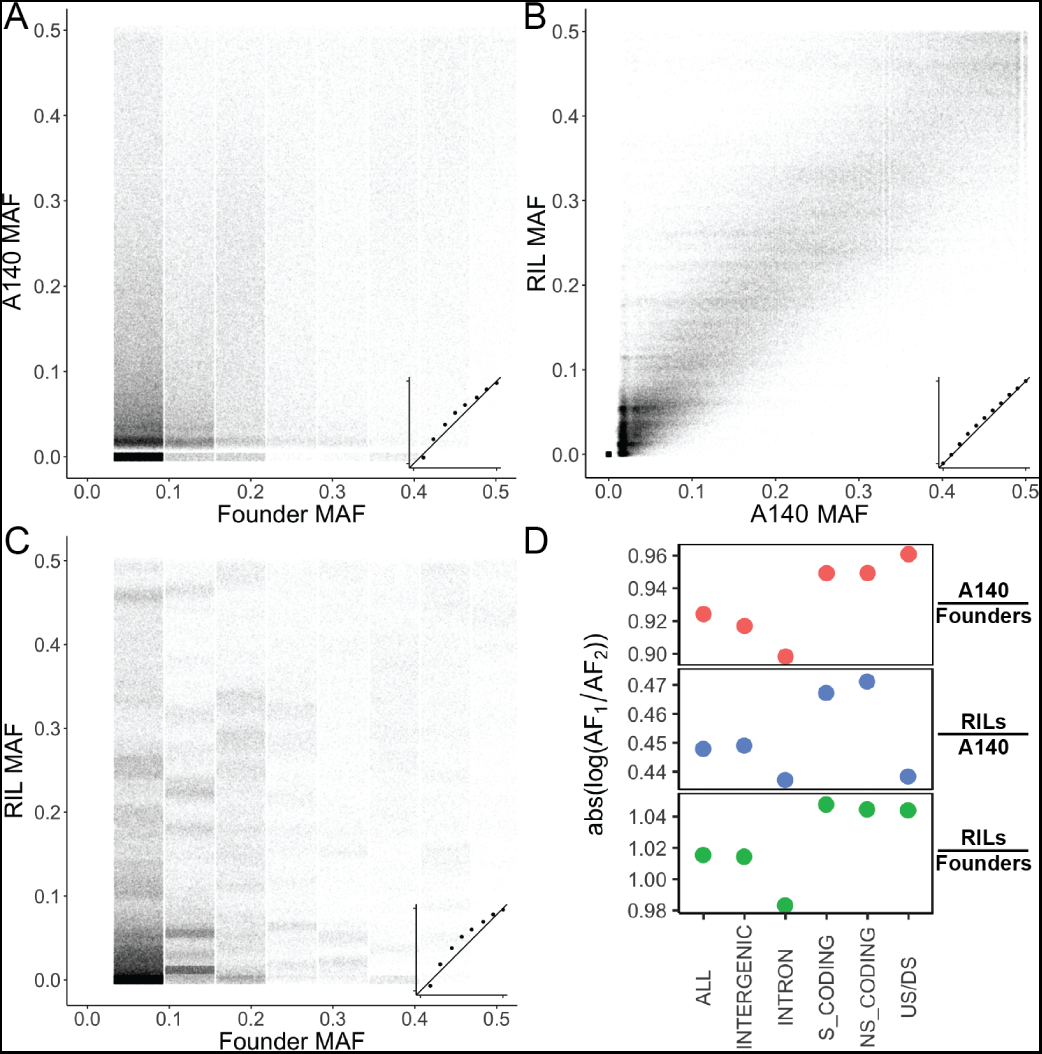
Minor allele frequency between founders and the outbred A140 population (**A**), A140 and RILs (inbreeding only for the A140 RILs, further adaptation then inbreeding for G50 RILs; **B**), and founders against all RILs (**C**). Insets show frequency quantiles. **D**. Change in allele frequency (absolute log ratios) for the same contrasts by functional class: intronic, synonymous and non-synonymous, putative regulatory variation (US/DS; ≤200bp from an annotated transcript or N2 pseudogene), or intergenic (none of the above). Points are mean values (diameter exceeds the standard errors).

Analysis of differentiation across variant functional classes showed large departures in frequency for coding variation (synonymous and non-synonymous) and the smallest for intronic variation (Figure 3D). Putative regulatory variation was most variable across experimental phases, being the most dynamic class during the funnel intercross and initial adaptation (founders to A140) but below the mean value for generations between the A140 and RILs. This pattern was observed across all of the sub-panels that make up the CeMEE (not shown), notably the A140 RILs which differ from the outbred A140 by only inbreeding, suggesting differential dominance of coding and regulatory variation (Wray 2007; Gruber *et al*. 2012). Without sequence data for the outbred CA50, GA50, GM50 or GT50 populations, we cannot assess the impact of inbreeding on the fixation of alleles more generally. These effects are expected to depend on reproductive mode and selection (Charlesworth and Wright 2001; Morran *et al*. 2009; Chelo and Teotónio 2013;*Chelo et al*. 2014; Kamran-Disfani and Agrawal 2014) and will be addressed in future work.

Local linkage disequilibrium, while non-uniform among chromosomes, decays relatively rapidly on average, approaching background levels by 0.5cM (F_2_ map scale) on average (Figure 4 and Figure S2). Disequilibrium between pairs of loci on different chromosomes is, as expected, very weak (0.99, 0.95 quantiles = 0.538, 0.051 within chromosomes versus 0.037, 0.022 across chromosomes), with the prominent exception of a single pair of loci on chromosomes II and III (*r*^2^ > 0.5 between II:2,284,322; tagging an intact MARINER5 transposon (WBTransposon00000128) that harbors an expressed miRNA in the N2 reference, and III:1,354,894-1,425,217; a broad region of mostly unannotated genes, against maximum interchromosomal values for all other pairs *r*^2^ ≤ 0.27). Alleles in repulsion phase are rare across these regions (*p* < 10^−70^, Fisher Exact Test), absent in the founders, and present in only 1 of 124 wild isolates surveyed with unambiguous variant calls in these regions (*Caenorhabditis elegans* Natural Diversity Resource). This suggests the presence of at least one two-locus incompatibility exposed by inbreeding or, perhaps more likely given the uncertainties of reference-based genotyping, a transposon-mediated II-III transposition polymorphism among founders. Three founders contribute the chromosome II non-reference haplotype, but extremely poor read mapping in this region for these and other isolates, consistent with high local divergence as well as potential structural variation, means our short read data are not informative in resolving these alternatives.

**Figure 4.**
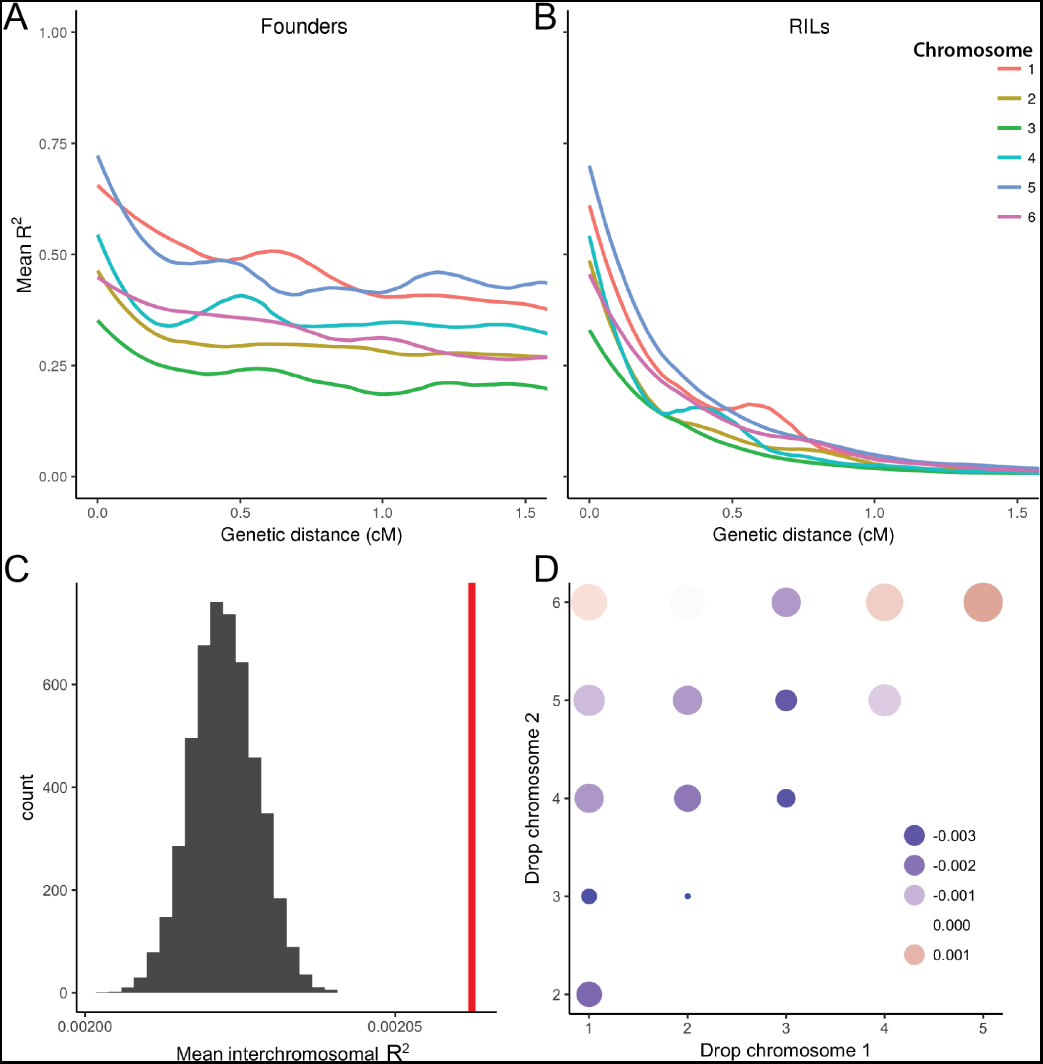
Linkage disequilibrium in founders (**A**) and all Ce-MEE RILs (**B**; F_2_ genetic map distance, LOESS fit to mean *r*^2^). **C.** Interchromosomal structure is weak but significant. Observed mean *r*^2^ across all chromosomes (red vertical bar) plotted against the null distribution from permutations randomizing lines across chromosomes (within sub-panels to exclude effects of population structure). **D.** Permutations dropping pairs of chromosomes implicate *X*-autosome interactions. Point size and color is scaled by enrichment over the null distribution (95% percentile), relative to the genome-wide mean value.

To better quantify the extent of subtle interchromosomal structure in the CeMEE we compared the observed correlations among chromosomes to values from permutations, shuffling lines within sub-panels, among chromosomes (Figure 4). The observed mean value for the genome, while extremely low, is highly significant (*p* < 2 × 10^−4^ from 5000 permutations), indicating the presence of extensive weak interactions. Further permutations dropping single or pairs of chromosomes showed that interactions between autosomes and the X chromosome contribute disproportionately.

### Founder haplotype blocks and genetic map expansion

The CeMEE panel is highly recombined and any simple, large-effect incompatibilities between founders are likely to have been purged. For example, a haplotype containing *peel-1* and *zeel-1*, a known incompatibility locus that segregates among the founders on the left arm of chromosome I (Seidel *et al*. 2008, 2011), is fixed in the RILs (Figure 5a). Cases such as this are best appreciated when the mosaic of founder haplotypes across the genome is inferred.

**Figure 5.**
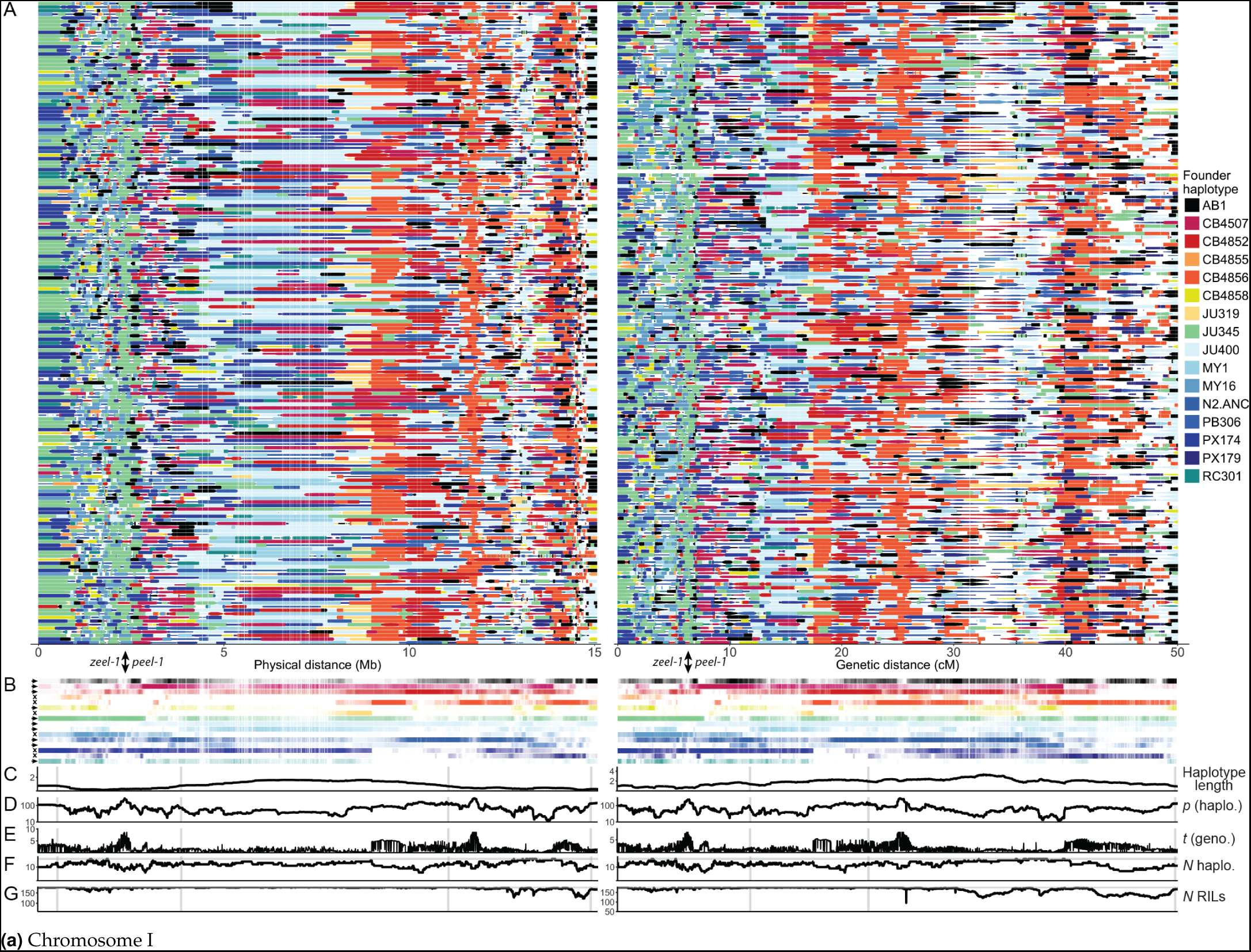
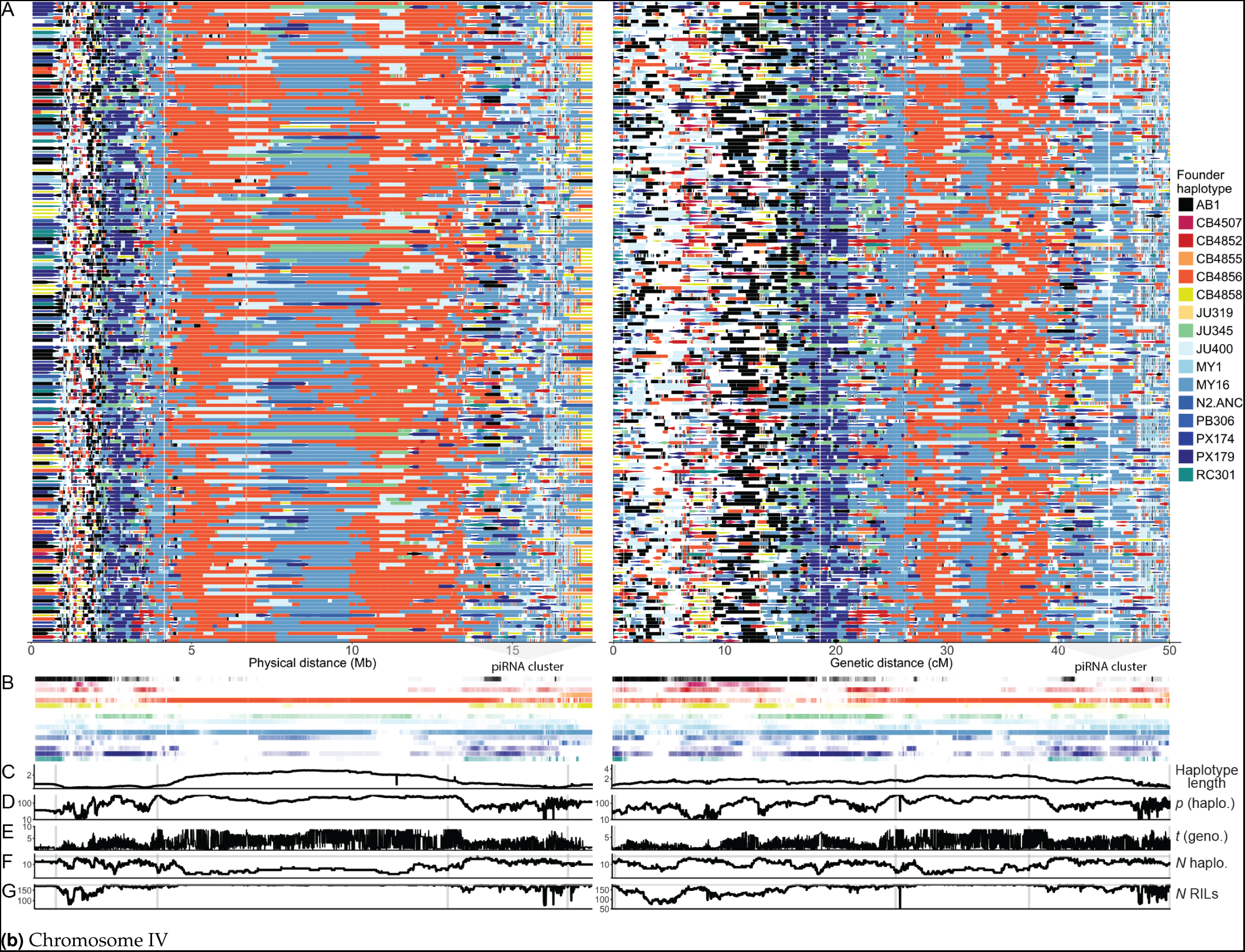
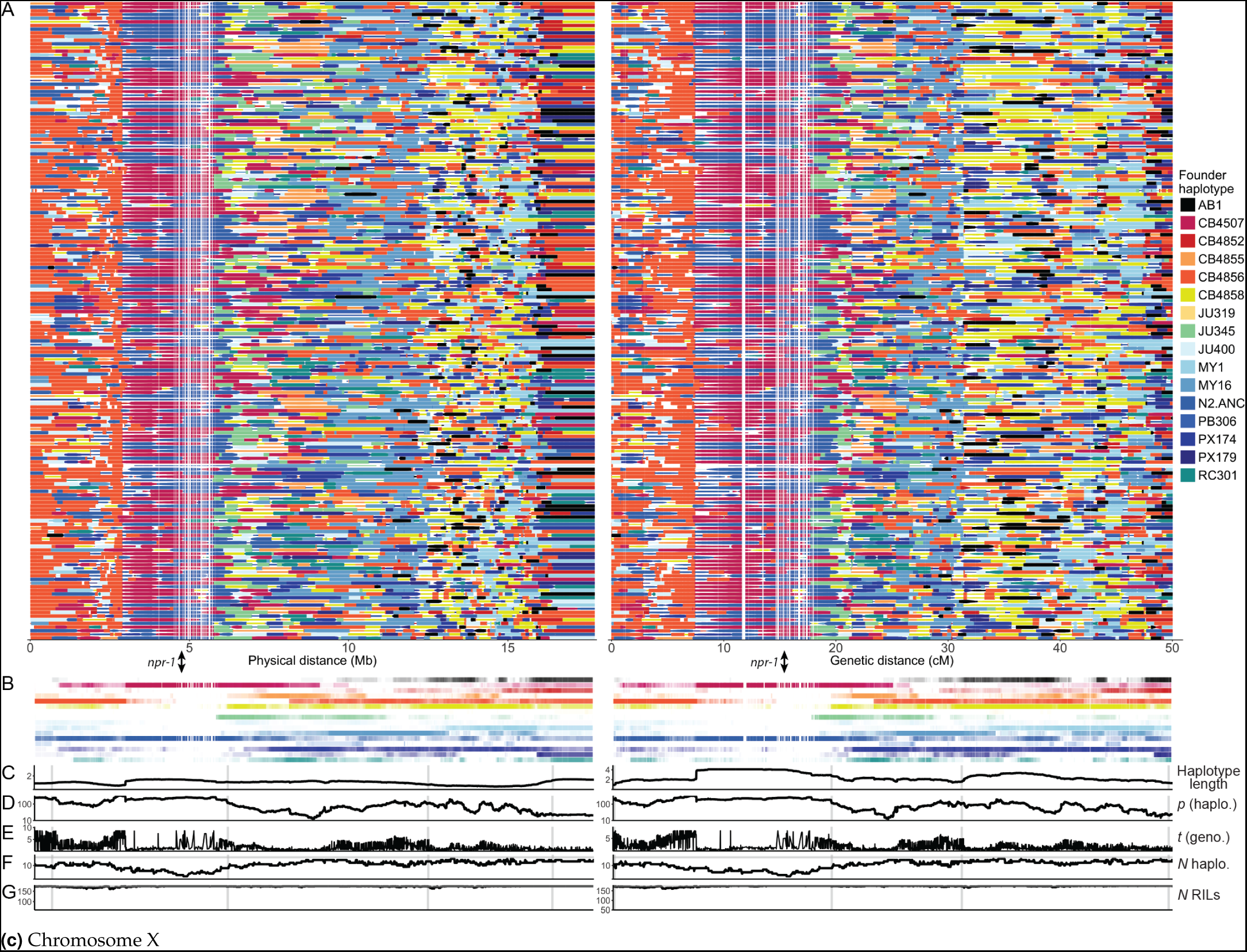
A140 RIL founder haplotype reconstruction and structure for chromosomes I, IV and X. **A**. Founder haplotypes reconstructed for the A140 RILs shown in physical and genetic distances. Each plotted point is a marker, with its size scaled by posterior probability (minimum 0.2). Founder contributions are summarized below in B. Loci discussed in the text are indicated: the *zeel-1/peel-1* incompatibility on the left arm of chromosome I (haplotype compatibility group, either experimentally tested in *Seidel et al*. (2008) or determined here from genotype data, is indicated below as an arrowhead for Bristol (N2) or an × for Hawaii (CB4856); extreme haplotype differentiation within a piRNA cluster on the right arm and tip of chromosome IV; and the fixation of N2/CB4507 haplotypes over a large region of the X chromosome left arm spanning *npr-1*, which has known pleiotropic effects on behavior and laboratory adaptation (de Bono and Bargmann 1998; Gloria-Soria and Azevedo 2008; McGrath *et al*. 2009; Andersen *et al*. 2014). **C-G** show summary statistics evaluated at 5Kb or 0.01cM resolution, with vertical scales for each metric fixed across chromosomes, and the positions of recombination rate boundaries inferred for the N2 × CB4856 RIAILs (Rockman and Kruglyak 2009) indicated with shaded bars. **C**. Haplotype length; mean length extending from the focal position. **D**. *p* (haplo.); test of reconstructed founder haplotype proportions, relative to expectation based on reconstruction frequency from G_150_ simulations (−*log*_10_(*p*) from a *χ*^2^ goodness-of-fit test). **E.** *t* (geno.); change in allele frequency from the founders (absolute value of Welch’s *t* statistic for founder vs. RIL genotype counts). **F.** *N* haplo.; the number of unique founder haplotypes detected at each position, with the maximum value of 16 indicated. **G**. *N* RILs; the number of RIL haplotypes reconstructed at each interval (> 0.2 posterior probability), with the maximum value of 178 indicated.

For each CeMEE RIL, founder haplotypes across the genome were reconstructed with the multiparent HMM framework RABBIT (Zheng *et al*. 2015), assigning 96.9% of markers to a single founder haplotype at posterior probability > 0.2 (84.2% > 0.5; median value across lines. Haplotype sharing in the 16 founders means that unambiguous assignment to a single founder is not always possible). For illustration purposes, a summary of reconstructed haplotypes for the A140 RILs on chromosomes I, IV and X are shown in Figure 5, at both physical and genetic scales to make the differences between these units plain. The observed recombination landscapes generally recapitulate those inferred from the N2/CB4856 cross (Rockman and Kruglyak 2009; Kaur and Rockman 2014; Bernstein and Rockman 2016), with recombination rate high in chromosome arms and low in centers. With the additional map expansion gained here (see below), we note that suppression of recombination is clearly strong, but not complete, within subtelomeric regions (see, for example, the exceptionally large right tip of chromosome X, spanning almost 2Mb, in Figure 5c).

Founder haplotype diversity among all CeMEE RILs remains high: the median number of founder haplotypes across reconstructed intervals is 12 (posterior probability > 0.5, haplotypes observed in > 1 RIL). Contributions clearly vary from equality, with lines most divergent from the reference (CB4856 and MY16) overrepresented and lines more similar to the reference underrepresented (with the exception of the large region of chromosome X, spanning *npr-1*, which is largely fixed for N2/CB4507 alleles (Figure 5c). To establish if these biases are merely technical, and establish expectations for reconstruction resolution in the presence of haplotype redundancy, we simulated genomes of varying pedigree length (up to 150 generations). As expected, reconstruction was hampered by increasing recombination, and by ambiguity between similar founders (Figure S3). Bias toward divergent haplotypes was not observed in the reconstruction simulations, however, suggesting the overrepresentation of CB4856 and MY16 may be due to selection, notably for long haplotypes across the central domain of chromosome IV (Figure 5b). Reconstruction completeness for the A140 RILs is generally in line with expectations for a pedigree of 150 generations. Clear exceptions are chromosome IV, where we recover more than expected under random sampling, and chromosome V, where we recover less. Haplotype lengths from simulated reconstructions showed we progressively underestimate recombination breakpoints due to imperfect resolution of small haplotypes (Figure S3).

The relationship between known generation and estimated realized map expansion from reconstruction simulations allows prediction of the number of effective generations of outcrossing within the CeMEE panel. Across the five sub-panels, mean autosomal generation ranges from 227 (GM monoecious RILs) to 284 (CA androdioecious lines), with a weighted average of 260 for the CeMEE as a whole (Figure S4). Estimated genetic map expansion is variable across chromosomes: IV appears to be exceptionally recombinant in all sub-panels with expansion more than twice that of chromosomes I-III, due largely to a high frequency, highly structured haplotype on the far right arm and tip (Figure 5b). This region spans one of the two large C. el*egans* piRNA clusters (Ruby *et al*. 2006), which encodes more than 15,000 piRNA transcripts, interspersed with active transposons and protein coding genes. Several trivial explanations for the unusual apparent expansion, such as elevated genotyping error rate or founder haplotype ambiguity, or distortions in the N2/CB4856 genetic map use to condition reconstruction probabilities, are not supported (data not shown), although the extent of large-scale structural variation among founders in this region (with the exception of CB4856, which does not show unusual levels of SNP or copy number variation) is unknown. Setting aside potential technical artifacts, the locus may represent a hitherto undetected recombination hotspot (whether through attraction, or suppression of observed recombination elsewhere on the chromosome), a site of rampant gene conversion, or the focus of early and sustained selection during the initial intercross phase (potentially epistatic in nature, see Neher and Shraiman (2009)). Earlier work proposed that evolution of this region may have involved a recombination rate modifier (through gene conversion) during the first 140 generations of experimental evolution in order to explain the observed excess haplotype diversity (see discussion and Figures S4 and S5 of Chelo and Teotónio (2013)). In contrast, chromosome V, which has been the focus of a recent large-scale selective sweep (Andersen *et al*. 2012), shows more variable expansion across sub-panels suggestive of ongoing selection (Figure S4).

### Population stratification

We examined additional genetic structure in the CeMEE RIL panel stemming from the inclusion of distinct sub-panels of RILs that vary in experimental evolution histories. In the context of QTL mapping, this genetic structure represents nuisance variation that can bias estimates of heritability if unknown factors covary with the trait of interest, structure that is causally associated with a trait, or non-causal structure due solely to population stratification.

To gauge the extent of population stratification we compared the results of supervised and unsupervised discriminant analysis of principal components (DAPC; Jombart (2008)), which partitions within and between group variation, using either known or inferred populations, based on linear combinations of principal components. By selection of discriminant functions that best predict known CeMEE sub-panel membership, it is clear that the varied evolutionary history has, unsurprisingly, generated significant genetic structure. The number of principal components selected by cross-validation that best predicts population membership is 40, which together explain 25% of the variance (though only a fraction of these components are significantly associated, considered singly or in pairs, see 3). Unsupervised DAPC, which infers groups based on variance minimization and model penalization criteria (*k*-means clustering, BIC), selected 5-8 clusters which best explain the data (Δ BIC < 1 over this range). These corresponded significantly with sub-panel identity (e.g., *p* = 0.036 at *k*=5, permutation test), although the rate of successful assignment was low (36% at *k*=5). This suggests that genetic structure within, as well as between sub-panels, is significant.

### Heritability and predictability of fitness-proximal traits

We measured two traits that are important components of fitness - the fertility and size of young adult hermaphrodites - and thus represent challenging case studies for mapping of complex traits in the panel (Poullet et al. 2016). The traits are correlated (Figure S1), and vary extensively in the CeMEE RILs: hermaphrodite fertility varies more than five-fold, size varies more than three-fold (Figure 7).

**Figure 6.**
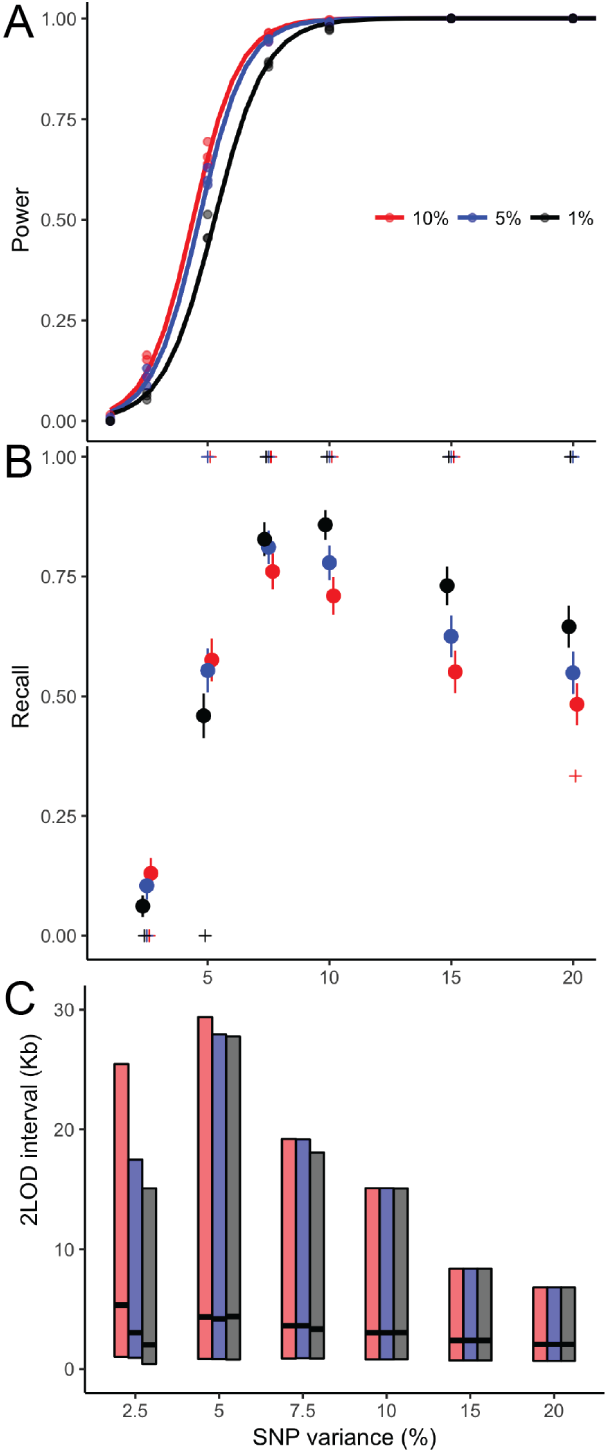
Additive QTL mapping simulations. Detection power (**A**), recall (**B**) and resolution (**C**; 2-LOD drop interval size for detected QTL) from single QTL simulations for the full mapping panel of 507 lines, as a function of detection threshold (significance at 0.01, 0.05 and 0.1) and phenotypic variance explained by the simulated QTL. Total heritability of simulated phenotypes is twice that of the focal QTL, with the polygenic contribution spread over 10, 100 or 1000 background markers (plotted in **A**, combined in **B** and **C**). In **B**, points are mean ± standard error. Recall declines with SNP variance at high levels as chance associations reach significance, although the median value (+ symbols) is 1.0 at 5% significance for variants that explain ≥ 7.5% of trait variance.

**Figure 7.**
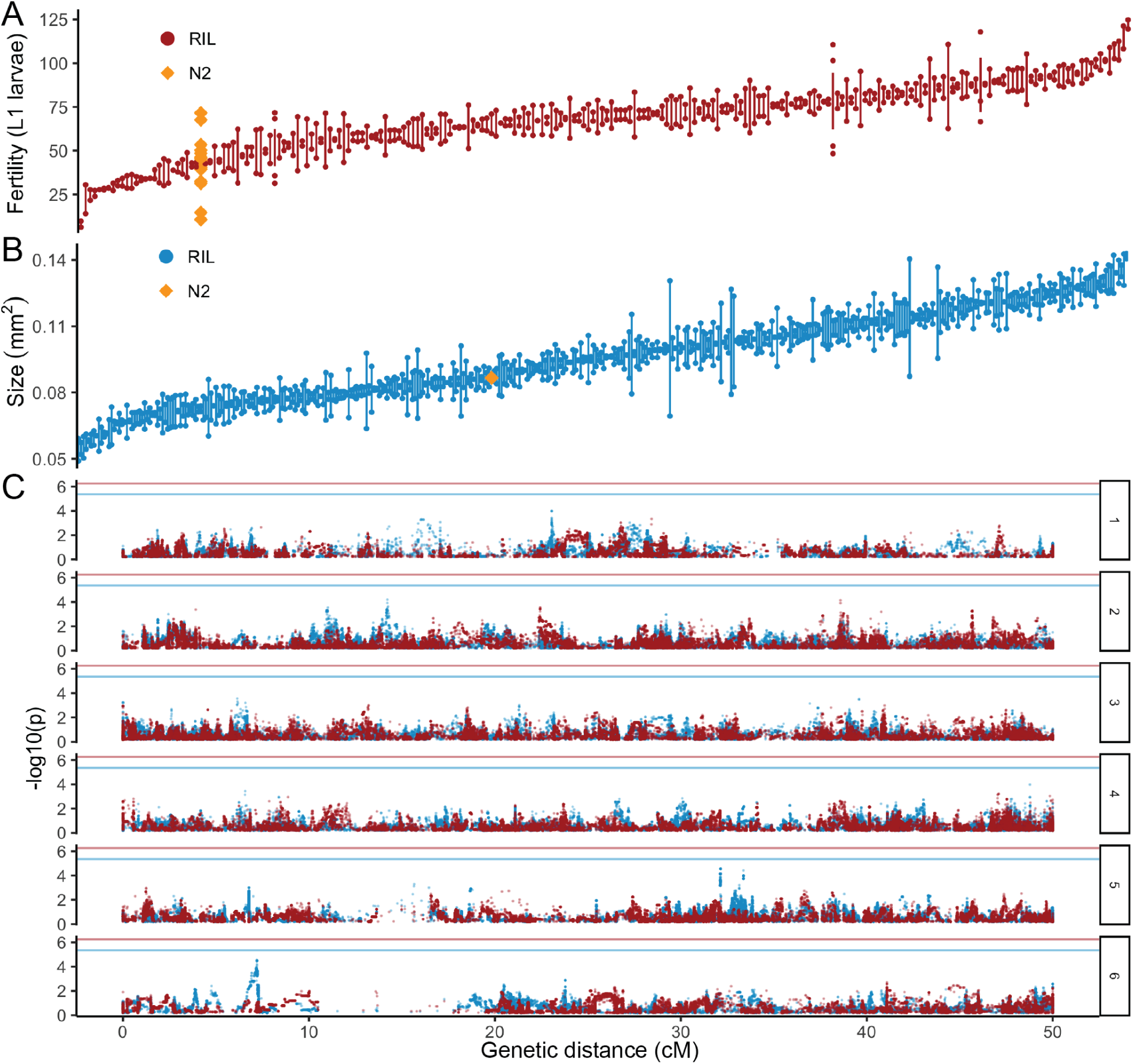
1D GWAS. **A-B**. Trait value distributions across RILs (replicate means; bars show data range or the standard error for samples with >2 replicates) and (**C**) single-SNP association results for fertility and adult body size (colors as above). Values for the reference N2 strain are shown for comparison. Note that values are raw replicate means on the original scale, and so include all sources of technical variation (unlike model coefficients used for mapping).

Under the uncommonly met assumptions of complete tagging of causal variation and uniform linkage, narrow sense heritability (*h*^2^) can be estimated from phenotypic and additive genetic covariances (Henderson 1975; Robinson 1991; Yang *et al*. 2010; Speed *et al*. 2012; de los Campos *et al*. 2015). Estimates, assuming appreciable heritability, are influenced by the extent to which markers reflect the genetic architecture of the trait in the population under study, and the method by which similarity is defined from them (reviewed in Speed and Balding (2015), and see Materials and Methods). Different covariance metrics can therefore provide useful information on the genetic basis of complex traits, such as partitioning chromosomal contributions, the frequency distribution of causal variants, and the proportion of epistatic variance, without the statistical limitations (and precision) of GWAS that attempt to explain phenotypic variance as the sum of individually significant additive marker effects. As emphasized by Speed and Balding, genomic heritability estimation is best viewed as a model-fitting exercise, the problem being to find the most appropriate measure of genetic similarity for the trait, population and genetic data in question, and the answer being the most likely estimate of the contribution of genetic variance to trait variance given the data.

### Repeatability, genomic heritability and prediction

While RIL repeatability (an upper bound on broad sense heritability, H^2^, under certain assumptions (Dohm 2002)) for both traits was relatively high − 0.76 for fertility and 0.80 for size - genomic heritability estimates for trait coefficients with a simple additive genetic similarity matrix based on the probability of allele sharing at all markers, equally weighted, were not significantly different to 0 (likelihood ratio test; not shown). This suggested that genome-wide genotypic similarity is poorly correlated with causal variation for these traits, potentially due to variable LD or epistatic cancellation. We thus examined alternative measures of genetic similarity to address the apparent lack of additive genomic heritability, comparing model predictive power (r^2^) by leave-one-out cross-validation (see Materials and Methods).

Heritability estimates and prediction accuracy are summarized in Table 1, comparing the simplest models – additive (A) only, or additive + additive-by-additive (A^2^) genetic covariance at the genome level – and the most predictive models for each trait. Given relatively high variance in relatedness, we are powered to detect large differences in additive heritabilities despite modest sample sizes for analysis of this kind, although the differences between individual models are generally minor. For fertility, with just 227 lines we have 50% power to reject *h*^2^ = 0 if *h*^2^ = 0.38, and >95% power at our estimate of *H*^2^ (at a significance level of 0.05), while for size, the corresponding values are 50% power at h^2^ = 0.35 and >99% power at repeatability (based on the best performing measure of additive similarity for each trait; Visscher et al. (2014)). Given the multiplicative scaling of epistatic similarity, low power is unavoidable.

**Table 1.** Genomic heritability estimates. Results are shown for additive (A) and additive-by-additive (A^2^) genetic similarity matrices (GSM), and for the most predictive model tested (if neither of the above), shown in bold. *α* is the scaling parameter from (Speed *et al*. 2012), which determines the effect size expectation for markers as a function of allele frequency, where 0 is unweighted and smaller values assign greater weight to rare alleles. Unconstrained REML estimates and standard deviations are shown for components that were >0 at convergence. *LR* is improvement over the null model (likelihood ratio). A+A^2^)_*rec*_ is additive and additive-byadditive similarity at the level of recombination rate domains (tips, arms and central domains).

| Trait | GSM | $\alpha$ | $r^2$ | $\hat{h}^2$ (SD) | LR |
| --- | --- | --- | --- | --- | --- |
| Size | A | -0.5 | 0.073 | A 0.58 (0.14) | 10.8 |
|  | **A+A<sup>2</sup>** | **-0.5** | **0.093** | **A 0.57 (0.15)** | **10.9** |
|  |  |  |  | **A<sup>2</sup> 0.21 (0.51)** |  |
| Fertility | A | 1 | 0.012 | A 0.24 (0.24) | 0.01 |
|  | A+A<sup>2</sup> | 1 | 0.029 | A 0.36 (0.21) | 2.67 |
|  |  |  |  | A<sup>2</sup> 1.24 (0.87) |  |
|  | **(A+A<sup>2</sup>)<sub>rec</sub>** | **1** | **0.064** | **A<sub>arm</sub> 0.44 (0.18)** | **6.98** |
|  |  |  |  | **A<sub>cen.</sub> 0.02 (0.07)** |  |

While phenotype prediction accuracy is generally poor, some broad trends are apparent in the ranking. Additive heritability based on LD-weighted markers was relatively high for size (0.58), though less so for fertility (0.24). In neither case was additive similarity alone the best predictor of phenotype, however. Nine of the top 10 models for fertility all incorporated epistasis in some form, with the best of these giving 57% improvement over the best additive model. For size, the advantage was less clear: three of the top four models included epistasis, though the performance differential between the best epistatic and additive models was only 3%.

Notably, partitioning of the genome based on recombination rate domains performed well for both traits, and was the preferred model for fertility. In general, model type was more influential on prediction than allele frequency scaling (*α*), however within models, negative values of *α* (rarer alleles having larger effects) were generally preferred for size, and positive for fertility, suggesting the frequency spectrum of causal variants for the two traits varies in the RILs.

### Effects of population stratification on heritability estimation

Given the stratified nature of the CeMEE panel, we tested for effects on heritability estimation in three ways. First, we estimated heritability for individual sub-panels (best additive models only). Although highly uncertain given the very small sample sizes, estimates were positive for two of the three sub-panels for adult body size and for both of two sub-panels tested (*n* > 50) for fertility, spanning the reported values for all lines.

Second, we estimated within sub-panel heritability by fitting within population means as covariates (best **A** and **A**+**A**^2^ models). For adult body size, where GA RILs are significantly larger than other panels, this reduced estimated heritability to 0.15 (A) and 0.38 (**A**+**A**^2^, with A^2^=0.30). Fertility, for which trait values vary only weakly with sub-panel, was largely unchanged at 0.45 (A) and (the unreasonably high, and uncertain) estimate of 1.44 (SD 0.75) for **A**+**A**^2^, with a dominant contribution from epistasis.

Third, we applied the method of Yang *et al*. (2011), developed for unrelated human populations, which compares the sum of heritabilities estimated for single chromosomes to that of a model fitting all chromosomes jointly. In the former case, genetic correlations across chromosomes due to population structure will result in 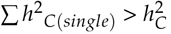, since the genotype of one chromosome will be predictive of that of others, while fitting all chromosomes jointly gives independent conditional estimates. The reasonable underlying assumptions are that structure is more significant between than within populations, and is not causally associated with phenotypic variance, although the latter might not hold for fitness-proximal traits. Comparing the sum of heritability estimates from samples of half the chromosomes (Σ *h*^2^/_2_) to that from all chromosomes (additive similarity only), results suggested stratification may contribute significantly to our estimates for size, with mean Σh^2^/_2_=0.72 (contributing 20% of the total given h^2^=0.60 for a joint chromosome model), but not for fertility (mean 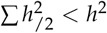 < *h*^2^). Fitting up to 80 principal components as covariates for size failed to bring this ratio to equality, but progressively eroded the heritability estimate (minimum 10% inflation for 80 PCs, h^2^=0.30), while fitting three DAPCs (based on the top 40 PCs) fully accounted for the difference (mean 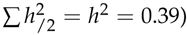. Notably, performing the same analysis within sub-panels, however, gave a similar level of ‘inflation’’ for size within the largest group of RILs (28%), suggesting that structure not associated with sub-panel is also influential.

The above analyses lead us to conclude that results presented in Table 1 for fertility are robust, while those for adult size are somewhat less so. The extent of inflation, however, is unlikely to be as severe as indicated by disjoint genome partitioning, and no covariates were fit for subsequent analyses.

### GWAS

#### QTL mapping power and precision

We first explored the characteristics of the CeMEE panel relevant to mapping quantitative traits. We carried out association tests by linear mixed effects model on simulated phenotypes, varying the effect size of causal variation and the degree of polygenicity (see Materials and Methods). The panel reaches 50% power for an allele explaining 0.047 of the phenotypic variance (permutation 5% significance threshold of *p* < 1.62 × 10^−6^), with recall (% true positives) greater than 50%, (Figure 6). When detected, the median QTL support inter-val (a drop in LOD of 2) spans < 10Kb for variants explaining >2.5% of trait variance. Given an average gene size of approximately 5Kb in *C. elegans* N2, including intergenic sequence, the CeMEE reaches sub-genic resolution for alleles of moderate effect (>10%), yielding high mapping precision (Figure 6). We note that our simulations are unbiased with respect to chromosomal location, while causal variation for many traits may be enriched on the highly recombinant arms, so these estimates are likely to be conservative.

#### 1D mapping of fertility and size

We carried out single marker genome-wide association tests by linear mixed effects model, controlling for genome-wide relatedness using the most predictive LD-weighted additive genetic similarity matrix for each trait (see above). Based on permutation thresholds, no single marker reached significance in either case (*α* = 0.1 thresholds = 4.38 ×10^−6^ and 5.57x10^−7^ for size and fertility, with minimum observed *p*-values of 2.8×10^−5^ and 7.23×10^−5^ respectively; Figure 7). For size, p-values were moderately inflated at the high end, with a number of regions approaching significance, but were strongly deflated for fertility, consistent with model misspecification. Results were largely independent of the method used to define similarity or, for fertility, whether correction for relatedness was applied at all (Figure S5). LD score regression, a related approach that explicitly assumes an infinitesimal architecture (Bulik-Sullivan *et al*. 2015), gave further support for extensive polygenicity with effects distributed across the genome (again, mostly clearly for fertility; Figure S6). Given significant heritabilities for both traits, and the results of GWAS simulations, the absence of individually significant associations suggests architectures comprising many variants with additive effects explaining <5% of the phenotypic variance.

#### 2D mapping of additive-by-additive interactions

Given suggestive evidence for epistasis from variance decomposition and a lack of individually significant additive effects by 1D mapping, we sought to identify interactions by explicitly testing pairs of markers. As summary statistics we retained the ANOVA interaction F statistic, as well as the sum of values for each marker across all tests for a chromosome pair (thresholded at F>0,8 and 16, the latter corresponding approximately to the most significant 1D associations seen). At a significance level of α=0.1 we detect four interactions (between seven loci) for fertility and two for size, with modest marginal additive effects (Figure 8; best single-locus statistics per pair ranging *p* = 9.1 × 10^−3^ − 9.9 × 10^−5^ for fertility and p = 1.1 × 10^−3^ − 9.9 × 10^−6^ for size). The variance explained by each pair, considered individually, is high: 12-15% for fertility and 7-8% for size, and a joint linear model explains 32% and 15% of the phenotypic variances.

**Figure 8.**
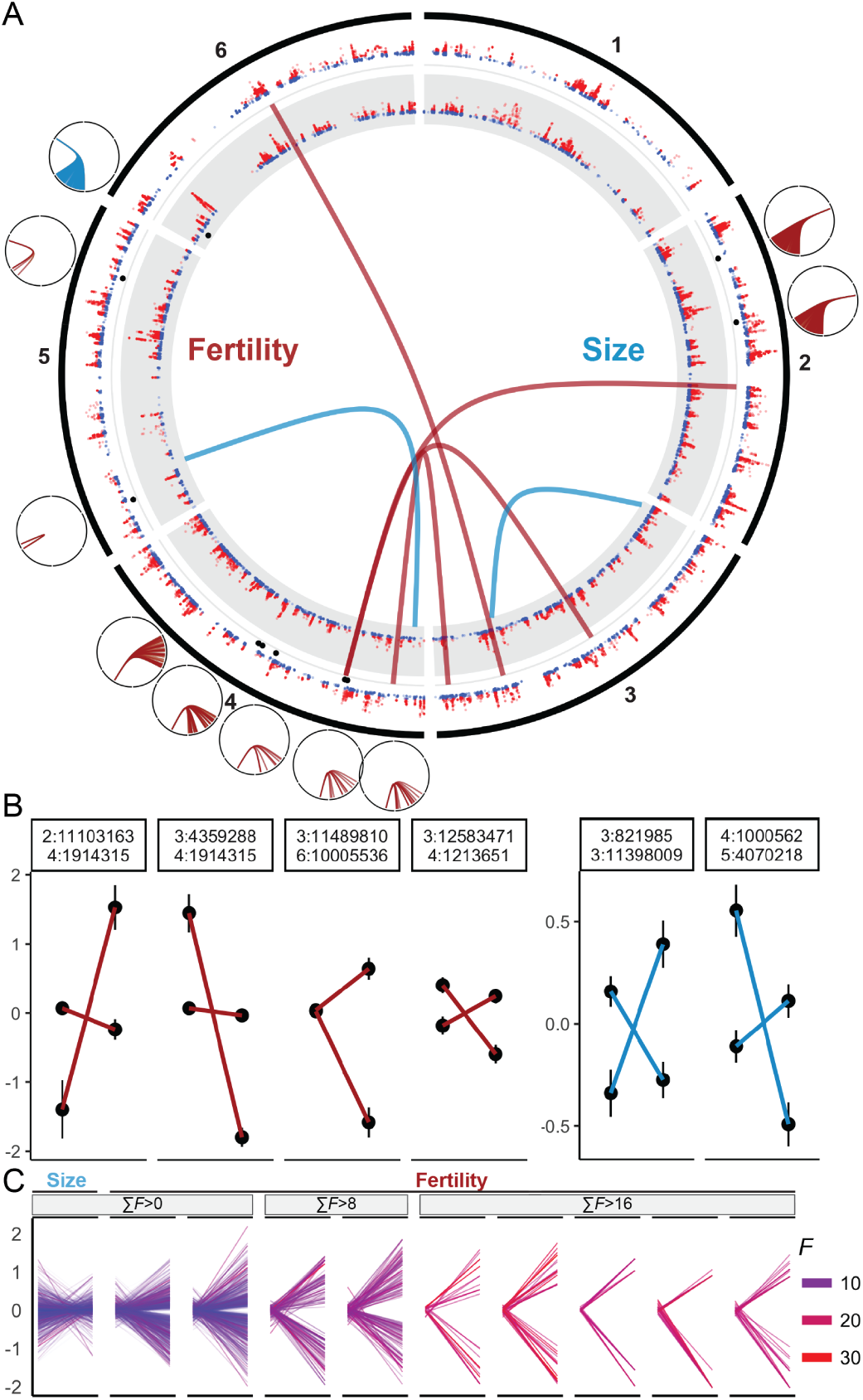
Strong sign epistasis and highly polygenic interactions contribute to trait variance. A. The distribution of significant interactions for fertility and size (genetic distance). Pairwise interactions are plotted over 1D GWAS test statistics (−log10(*p*) > 1) for each trait. Markers with a significant excess of summed interactions for a given chromosome pair are indicated with black points, and the chromosome identities and locations of interacting loci are shown as smaller plots at their approximate positions. 2D sum tests are directed interactions between a single focal marker, and all other markers on one other chromosome, with the sum of interaction scores reaching significance (*α* = 0.1) under a null permutation model. Note interactions between chromosome V:3,145,783 and 16 loci on the right tip of chromosome IV are clustered over a physical interval of 0.44Mb (in weak LD) and appear as a single link at this resolution. B. Genotype class trait means (± SE) for significant pairs (fertility in red, size in blue). C. Genotype class trait means for all individual pairs that contribute to significant summed interactions, at each of the three evaluated *F* statistic thresholds (interactions significant at F>0 are filtered to F>2 for plotting). Line color and intensity is scaled by F for each constituent interaction. Strong sign epistasis (including weak reciprocal sign epistasis) is the prevalent epistatic mode.

By summing interaction scores in 1-dimensional space to test for polygenic epistasis, we detect 10 unique markers with excess interchromosomal interactions for 3 chromosome pairs for fertility (a=0.1, across all three minimum *F* threshold classes), and one for size (at F>0; Figure 8). Only one of these sites also reaches significance in single pair tests: position 1,914,315 on chromosome IV, which is involved in individually significant interactions of opposite effect with chromosome II and III for fertility, and, remarkably, has at least one interaction of weak to moderate effect (10^−5^ < *p* < 10^−4^) with all other chromosomes. A flanking marker in modest linkage disequilibrium (IV:1,894,021, r^2^ = 0.31) also shows a significant excess of interaction scores with chromosome III that do not appear to be driven solely by LD: 6/12 interactions (F > 16 for IV:1,894,021) are shared with IV:1,914,315, and among all 26 interactions involving these two sites (F > 16 for either), interactions statistics are uncorrelated (*r* = -0.15, *p* = 0.49). Nevertheless, experiment will be required to test these loci independently.

IV:1,914,315 is found within an intron of *egl-18* (encoding a GATA transcription factor), while IV:1,894,021 falls within the large intergenic region between *egl-18* and *egl-4* (encoding a cyclic-GMP-dependent protein kinase thought to act in the TGF-beta pathway), both of which vary in coding and UTR sequence among founders, and have numerous known phenotypes from classical induced mutations and RNAi spanning the gamut of behavior, development and reproduction. Their eponymous phenotype, egg-laying abnormal (Egl), is retention of oocytes and embryos, a phenotype selected during experimental evolution in which embryos were extracted each generation by bleaching (Poullet *et al*. 2016).

## Conclusions

We have described the generation, characterization and application of the first multiparental mapping panel for the model organism *C. elegans*. Drawing on effectively 260 generations of moderate population sizes and predominant outcrossing during laboratory culture, full reference-based genome sequencing of the 16 inbred wild founders, and dense genotyping of the RILs, the CeMEE panel yields gene level mapping resolution for alleles of 5% effect or greater. For traits such as gene expression, for which the proportion of variance explained by local variation is typically upwards of 20% (e.g., Brem and Kruglyak (2005); Rockman *et al*. (2010); King *et al*. (2014), the majority of QTL intervals will dissect single genes.

While reference-based genotyping will remain a necessity for some time yet, it leaves the contribution of certain classes of genetic variation uncertain, and can hamper variant calling due to mapping bias and erroneous alignments at copy number variants. The genome of only one wild-isolate, the Hawaiian CB4856, has been assembled *de novo* to a high standard, revealing extensive divergence (Thompson *et al*. 2015). The ultimate goal of full genomes for all founders will yield both better accuracy in calculating genetic similarity, and ability to measure the phenotypic effects of this recalcitrant variation. Similarly undetermined, given RIL genotyping by mostly low coverage sequencing, is the extent and fate of novel mutations during experimental evolution. With a mutation rate of around 1/genome/generation for SNPs, and more for multinucleotide mutations and copy number variation (Denver *et al*. 2004a,b; Seyfert *et al*. 2008; Denver *et al*. 2010; Phillips *et al*. 2009; Lipinski *et al*. 2011; Meier *et al*. 2014), the contribution of new mutations to trait variation in the RILs may well be non-negligible. Theory suggests that fixation of adaptive mutations should not be significant during experimental evolution (Hill 1982; Caballero and Santiago 1995; Matuszewski *et al*. 2015), but empirical evidence is mixed (Estes 2004; Estes *et al*. 2011; Denver *et al*. 2010; Chelo *et al*. 2013). Both of these factors would erode phenotype prediction accuracy, which, theoretically, should converge on *H^2^* given perfect genotyping of all causal variation and appropriate description of genetic covariance (de los Campos *et al*. 2015).

The native androdioecious mating system of *C. elegans* and the ability to archive strains indefinitely confer significant advantages to further use, bestowing almost microbial powers on a metazoan model. For one, the preservation of intermediate outbred populations means that the CeMEE is readily extensible, limited only by effective population sizes. However, RIL panels have several potential shortcomings. First, despite inbreeding during RIL construction, a nagging concern in use of RIL panels is residual heterozygosity (Barrière *et al*. 2009; Chelo *et al*. 2014), and the possibility of further evolution of genotypes and phenotypes subsequent to characterization. While heterozygosity appears to be at a low level in the CeMEE RILs, on average, it is not absent (see Materials and Methods). Importantly, however, given that lines are in stasis the opportunity for segregation during further use is both limited and known. A second concern is the possibility of inbreeding depression, particularly for fitness-proximal traits. This is a concern for predominantly outcrossing organisms (Barrière *et al*. 2009; Philip *et al*. 2011; King *et al*. 2012; Chelo *et al*. 2014), but it is also applicable to multiparental experimental evolution of *C. elegans*. As mentioned in the introduction, at least during the initial stage of laboratory adaptation, excess heterozygosity may have been maintained by epistatic overdominant selection, and closely linked recessive deleterious alleles in repulsion could be maintained by balancing selection during inbreeding (Chelo *et al*. 2013, 2014). Assaying the *F_1_* progeny of nested crosses among RILs may be a useful approach to estimate (or avoid) the effects of inbreeding depression (Long et al. 2014).

Using subsets of the CeMEE panel, we outlined the genetics of two traits associated with fitness. Fertility, as defined here by the experimental evolution protocol employed, is correlated with hermaphrodite body size at the time of reproduction (Poullet et al. 2016). For both traits, and size in particular, additive genomic heritability based on LD-weighted similarity explained a significant fraction of *H*^2^, although heritability estimates were generally higher with the inclusion of epistatic similarity. This is consistent with a polygenic architecture with additive effects below the detection limit, whether solely additive, or due to weak or opposing effects of multiple interactions. Variance in fitness-related traits, in particular, may be maintained despite consistent selection on additive variation through a number of processes, including stabilizing selection under a stable environment (Whitlock *et al*. 1995; Wolf *et al*. 2000; Barton and Keightley 2002; Phillips 2008; Hemani *et al*. 2013). Results from variance decomposition, phenotype prediction and interaction tests are all consistent with this prediction: phenotypic variance remains high, and we find support for epistasis for both traits. Notably for fertility, which is expected to be well aligned with fitness under the experimental evolution scheme, strong interactions among four pairs of alleles with weak marginal main effects jointly explain almost a third of the phenotypic variance. All six interactions detected for fertility and size are instances of sign epistasis, where the directional effect of one allele is reversed in the presence of another. Five of these represent the extreme form, reciprocal sign epistasis (the reversal is, to some extent at least, symmetric; Poelwijk *et al*. (2011)). Sign epistasis, in particular, has important implications for a population’s capacity to adapt, by creating rugged fitness landscapes and constraining exploration of them (Weinreich *et al*. 2005, 2013), and for the repeatability of evolution, since the outcome of selection on the marginal additive effects of interacting alleles will be determined by their relative frequencies (Wright 1932;*Whitlock et al*. 1995; Phillips *et al*. 2000). Our tests for excess interactions among individually non-significant marker pairs additionally revealed a number of cases of highly polygenic epistasis, again, mostly for fertility. While tests of this type have the unsatisfying property of leaving the identities of the interacting partners uncertain, they have the potential to combat the loss of power that comes with explicit 2-dimensional testing (Crawford *et al*. 2016).

Fertility and body size at reproduction show broad-sense heritabilities that are relatively high for fitness-proximal traits (Lynch and Walsh 1998). This high heritability is likely a consequence of novel genetic variation created in the multiparental cross and realignment of selection to novel laboratory environments. While all mapping panels are synthetic systems, the mixing of natural variation and experimental evolution represents a perturbation that may have some parallels, for example, with that of a simultaneous founder event and environmental change, which can reveal novel incompatibilities and promote further differentiation (Cheverud and Routman 1996; Wolf *et al*. 2000). In this context, it will be useful to determine the directional effects of epistasis on the genotype-phenotype map during further evolution, as a function of recombination, a task for which the CeMEE is well suited. As in other systems such as Arabidopsis, where similar resources exist (Weigel 2012) and epistasis for fitness-related traits has been found (e.g., Malmberg *et al*. (2005); Simon *et al*. (2008)), it will also be important to begin a comprehensive comparison of QTL for fitness traits in the CeMEE and natural populations - where linked selection coupled with predominant selfing and meta-population dynamics have generated limited, structured genetic diversity (Andersen *et al*. 2012; Rockman *et al*. 2010; Cutter 2015) – and also with mutational variances obtained in mutation accumulation experiments (Baer *et al*. 2005; Baer 2008; Joyner-Matos *et al*. 2009). Such comparisons have the potential to provide significant insights into how the distributions of QTL effects and frequencies are shaped in natural populations.

## Acknowledgements

We thank J. Costa, R. Costa, C. Goy, F. Melo, H. Mestre, V. Pereira, and A. Silva for support with worm handling, sample preparation, and data acquisition; E. Andersen, M.-A. Félix, P.C. Phillips, S. Proulx, D. Speed and C. Zheng for helpful discussion.

## Author contributions

CeMEE panel derivation: S.C., B.A., H.T.; sequencing and genotyping: A. P.-Q., D.R., I.C. P.A., L.N., M.R.; phenotyping: I.C., B. A, A.C.; analysis: L.N., I.C., T.G., A.D, B.S.; manuscript: L.N., M.R., H.T.

## Supplementary figures

**Figure S1.**
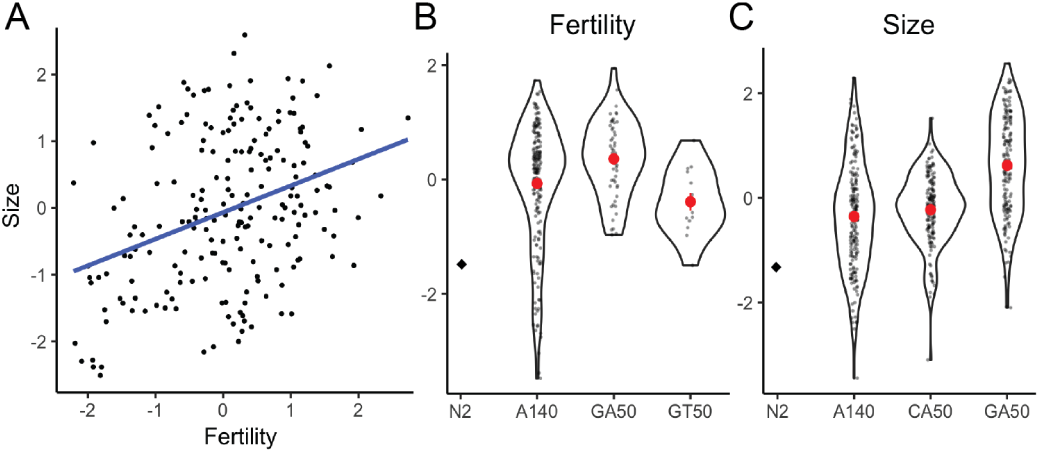
S1. Trait correlations and evolution. A. Fertility and size are correlated traits (Spearman’s *ρ* = 0.318, *p* < 5 × 10^−6^ for 202 RILs with data for both traits). **B-C**. Trait distributions within sub-panels (density plots and mean ± SE for centered and scaled model coefficients). The GA50 RILs are significantly larger and more fertile than A6140 RILs.

**Figure S2.**
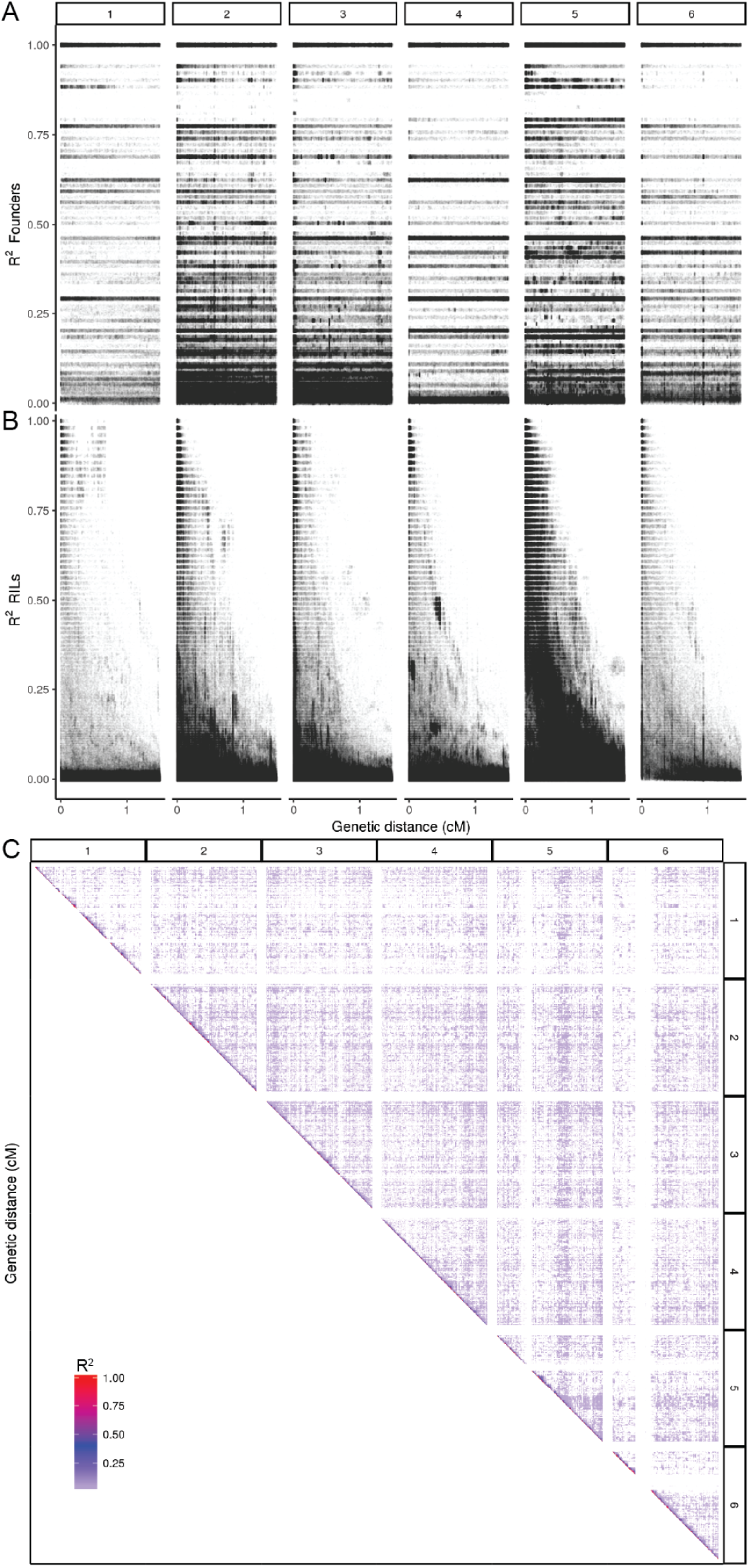
Local linkage disequilibrium in founders **A** and CeMEE RILs **B**, and across RIL genomes (**C**; *r*^2^ thresholded to >0.01).

**Figure S3.**
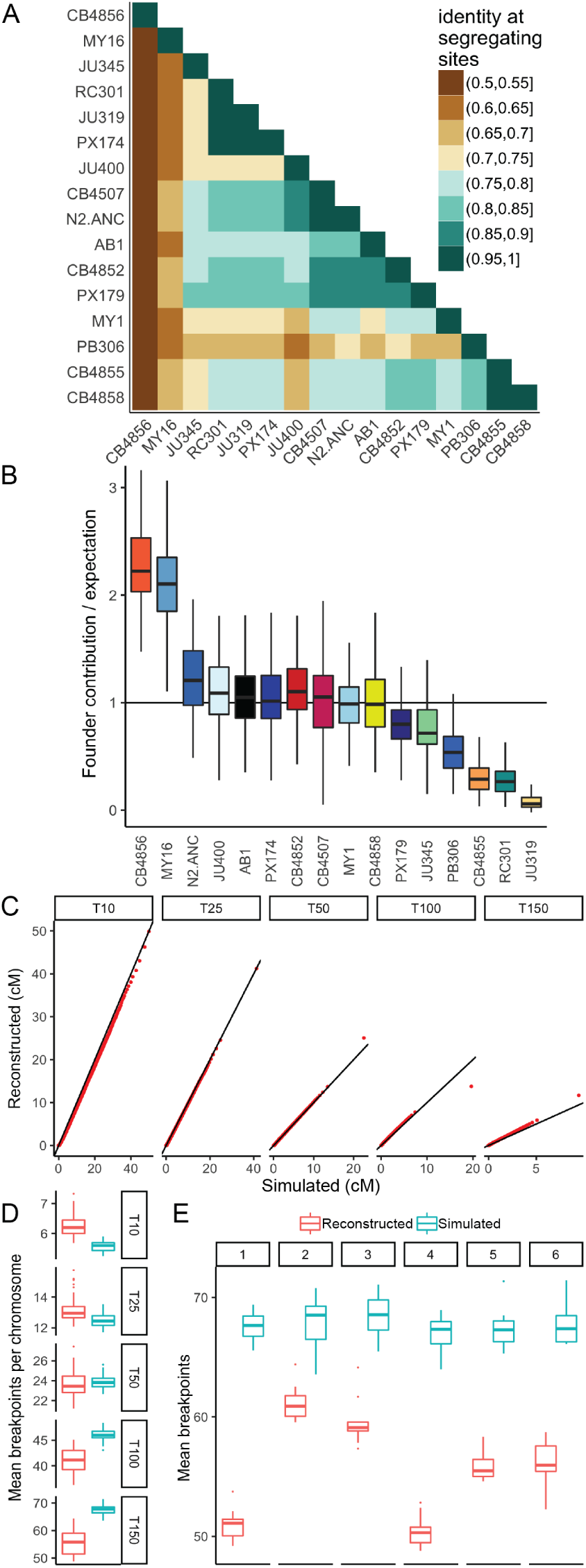
Summary of haplotype reconstruction. **A.** Genetic similarity among founders. **B.** Founder contributions in the CeMEE lines, relative to expectation from reconstruction of simulated recombinant genomes, accounting for bias based on haplotype uniqueness. Boxplots show median (bar), interquartile range (box) and 1.5 × the data range (whiskers). **C.** Haplotype length quantile-quantile plots for known and reconstructed simulations. **D.** The number of breakpoints per chromosome per line across simulation generation. **E.** The number of breakpoints reconstructed by chromosome. Haplotype uniqueness varies across chromosomes: the ability to reconstruct is poorest for chromosomes I and IV.

**Figure S4.**
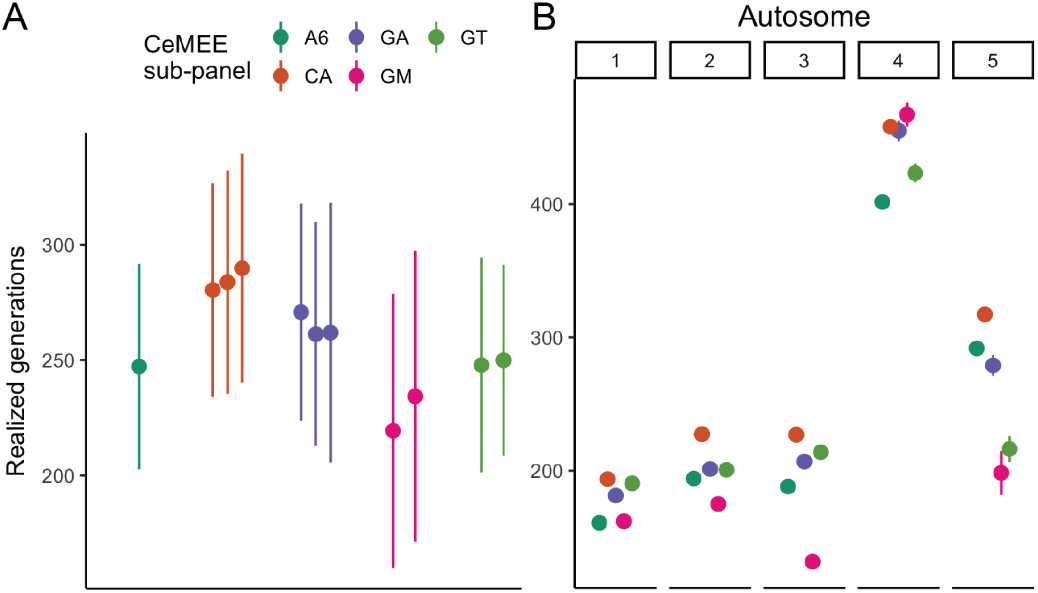
The number of generations of outcrossing for each CeMEE sub-panel (**A**) and chromosome (**B**) predicted from the maximium likelihood estimate of realized map expansion (Zheng *et al*. 2014, 2015).

**Figure S5.**
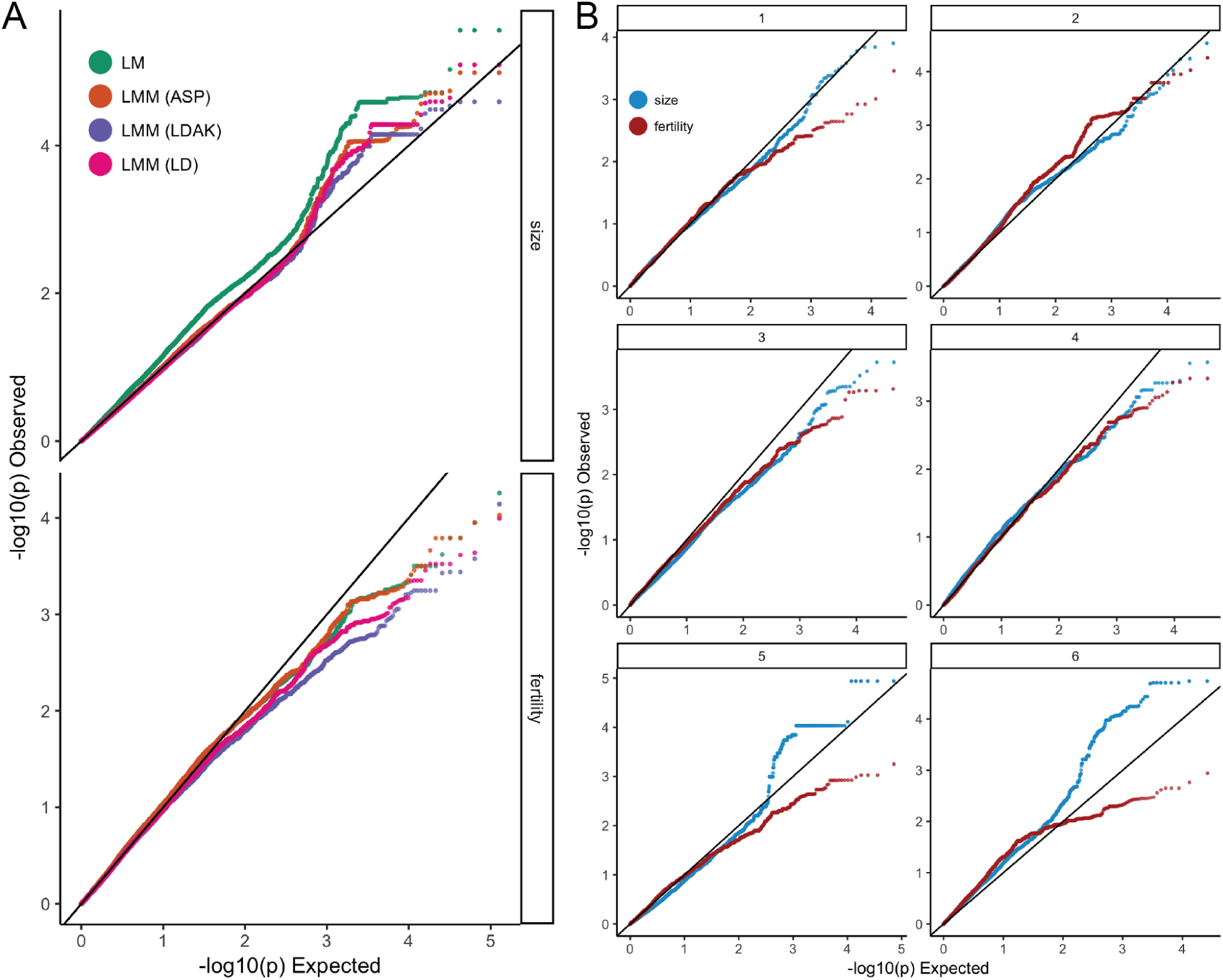
*p*-value quantile-quantile plots genome-wide (**A**), comparing the effects of relatedness corrections (where LM is linear model; LMM (ASP) is linear mixed model with relatedness based on allele sharing probability (all markers, equally weighted); LMM (LDAK) is the best performing LD-weighted similarity for each trait; LMM (LD) is based on markers pruned by local LD, but unweighted), and by chromosome (**B**), for the best LD-weighted similarity for each trait. While strong, spurious inflation is seen for size without polygenic correction (**A**), this is not seen for fertility, likely due the greater heterogeneity of trait values among sub-panels for size. Notably, deflation is seen for fertility for all models, although LD weighting introduces the strongest penalty, which may indicate a relationship between low LD and causal variation for this trait.

**Figure S6.**
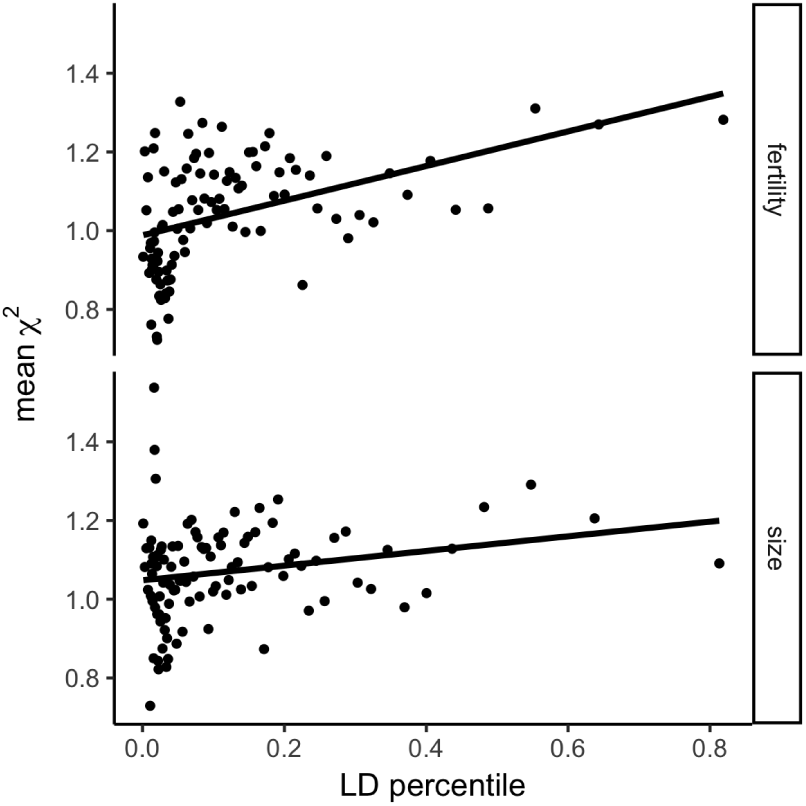
Fitness-proximal traits are polygenic. Regression of association statistics (mean value of χ^2^ percentiles) on marker LD weightings (mean of *w* percentiles, Speed *et al*. (2012)) for fertility and size (after Bulik-Sullivan *et al*. (2015)). While there is a significant positive relationship between trait association and the amount of variation tagged by markers, fertility shows much stronger evidence of polygenicity (slope=0.44, *p* = 2.7 × 10^−6^, versus slope=0.19, p = 0.029 for size).

